# Semi-soft Clustering of Single Cell Data

**DOI:** 10.1101/285056

**Authors:** Lingxue Zhu, Jing Lei, Lambertus Klei, Bernie Devlin, Kathryn Roeder

**Affiliations:** Department of Statistics and Data Science, Carnegie Mellon University; Department of Psychiatry, University of Pittsburgh School of Medicine; Department of Computational Biology, Carnegie Mellon University

**Keywords:** Single-cell RNA-seq, Soft clustering

## Abstract

Motivated by the dynamics of development, in which cells of recognizable types, or pure cell types, transition into other types over time, we propose a method of semi-soft clustering that can classify both pure and intermediate cell types from data on gene expression from individual cells. Called SOUP, for Semi-sOft clUstering with Pure cells, this novel algorithm reveals the clustering structure for both pure cells and transitional cells with soft memberships. SOUP involves a two-step process: identify the set of pure cells and then estimate a membership matrix. To find pure cells, SOUP uses the special block structure in the expression similarity matrix. Once pure cells are identified, they provide the key information from which the membership matrix can be computed. By modeling cells as a continuous mixture of *K* discrete types we obtain more parsimonious results than obtained with standard clustering algorithms. Moreover, using soft membership estimates of cell type cluster centers leads to better estimates of developmental trajectories. The strong performance of SOUP is documented via simulation studies, which show its robustness to violations of modeling assumptions. The advantages of SOUP are illustrated by analyses of two independent data sets of gene expression from a large number of cells from fetal brain.

---

Development often involves pluripotent cells transitioning into other cell types, sometimes in a series of stages. For example, early in development of the cerebral cortex (Kowalczyk et al., 2009), one progression begins with neuroepithelial cells differentiating to apical progenitors, which can develop into basal progenitors, which will transition to neurons. Moreover, there are diverse classes of neurons, some arising from distinct types of progenitor cells (Jones, 2009; Nadarajah et al., 2003). By the human mid-fetal period there are myriad cell types and the foundations of typical and atypical neurodevelopment are already established (Silbereis et al., 2016). While the challenges for neurobiology in this setting are obvious, some of them could be alleviated by statistical methods that permit cells to be classified into pure or transitional types. We will develop such a method here. Similar scenarios arise with the development of bone-marrow derived immune cells, cancer cells and disease cells (Keren-Shaul et al., 2017), hence we envision broad applicability of the proposed modeling tools.

Different types of cells will have different transcriptomes or gene expression profiles (Silbereis et al., 2016). Thus, they can be identified by these profiles (Darmanis et al., 2015), especially by expression of certain genes that tend to have cell-specific expression (marker genes). Characterization of these profiles has recently been facilitated by single cell RNA sequencing (scRNA-seq) techniques (Tang et al., 2009; Ramsköld et al., 2012), which seek to quantify expression for all genes in the genome. For single cells, the number of possible sequence reads is limited and therefore the data can be noisy. Nonetheless, cells of the same and different cell types can be successfully clustered using these data (Camp et al., 2015; Darmanis et al., 2015; Baron et al., 2016; Zeisel et al., 2015; Tasic et al., 2016).

What is missing from the clustering toolbox is a method that recognizes development, with both pure type and transitional cells. In this paper, we develop an efficient algorithm for Semi-sOft clUstering with Pure cells (SOUP). SOUP intelligently recovers the set of pure cells by exploiting the block structures in cell-cell similarity matrix, and also estimates the soft memberships for transitional cells. We also incorporate a gene selection procedure to identify the informative genes for clustering. This selection procedure is shown to retain fine-scaled clustering structures in the data, and substantially enhances clustering accuracy. Incorporating soft clustering results into methods that estimate developmental trajectories yields less biased estimates developmental courses.

We first document the performance of SOUP via extensive simulations. These show that SOUP performs well in a wide range of contexts, it is superior to natural competitors for soft clustering and it compares quite well, if not better, than other clustering methods in settings ideal for hard clustering. Next, we apply it to two single cell data sets from fetal development of the prefrontal cortex of the human brain. In both settings SOUP produces results congruent with known features of fetal development.

## Results

### Model Overview

Suppose we observe the expression levels of *n* cells measured on *p* genes, and let *X* ∈ ℝ^*n*×*p*^ be the cell-by-gene expression matrix. Consider the problem of semi-soft clustering, where we expect the existence of both (i) pure cells, each belonging to a single cluster and requiring a hard cluster assignment, as well as (ii) mixed cells (transitional cells) that are transitioning between two or more cell types, and hence should obtain soft assignments. With *K* distinct cell types, to represent the soft membership, let Θ ∈ ℝ_+_^*n*×*K*^ be a nonnegative membership matrix. Each row of the membership matrix, Θ_*i*_:= (*θ_i_*_1_, ⋯, *θ_iK_*), contains nonnegative numbers that sum to one, representing the proportions of cell *i* in *K* clusters. In particular, a pure cell in type *k* has *θ_ik_* = 1 and zeros elsewhere.

Let *C* ∈ ℝ^*p*×*K*^ denote the cluster centers, which represent the expected gene expression for each pure cell type. When a cell is developing or transitioning from one category to another, it may exhibit properties of both subcategories, which is naturally viewed as a combination of the two cluster centers. Weights in the membership matrix reflect the stage (early or late) of the transition. Here we formulate a simple probability model that is convenient for analysis and highly robust to expected violations of the assumptions. Let

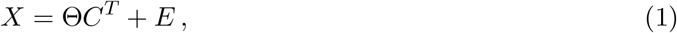

where *E* ∈ ℝ^*n*×*p*^ is a zero-mean noise matrix with 𝔼(*EE^T^*) = *σ*^2^*I*. It follows directly that the cell-cell similarity matrix takes a convenient from:

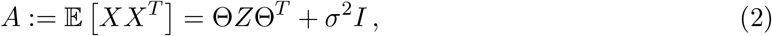

where *Z* = *C^T^ C* ∈ ℝ^*K*×*K*^ represents the association among different cell types.

In practice, many genes will not follow the developmental trajectory described by eq.(1); however, it is expected that the expression of many marker genes and other highly informative genes will transition smoothly between cluster centers during development (see, for example the genes featured in Trapnell et al. (2014)). In particular, one can empirically check the plausibility of eq.(1) for marker genes; see the “Case Studies” section below for details. Moreover, because SOUP’s inferences are based on the empirical cell-cell similarity matrix 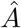, it is sufficient that 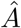 approximately follows the form specified in eq.(2), a weaker assumption than eq.(1). Indeed similar assumptions are implicit in many algorithms that estimate developmental trajectories (Bendall et al., 2014; Shin et al., 2015; Ji & Ji, 2016; Street et al., 2018). Gene expression is also likely to have non-constant variance, depending on gene and cell type. However, our pure cell search algorithm does not depend on the diagonal entries of *A*, and our estimate of Θ is based on spectral decomposition of *A*, so the method remains robust to moderate fluctuation of diagonal entries of *A* unless the magnitude of noise is unrealistically large.

As a graphical illustration of the SOUP model, we simulate an example with a developmental trajectory of type1 → type2 → type3. A fraction of the genes were chosen to have differential expression across cell types, and of these a fraction change nonlinearly between cell types (Figure 1a). Regardless of the violations of eq.(1), the cells depict a smooth transition between cell types (Figure 1b).

**Figure 1:**
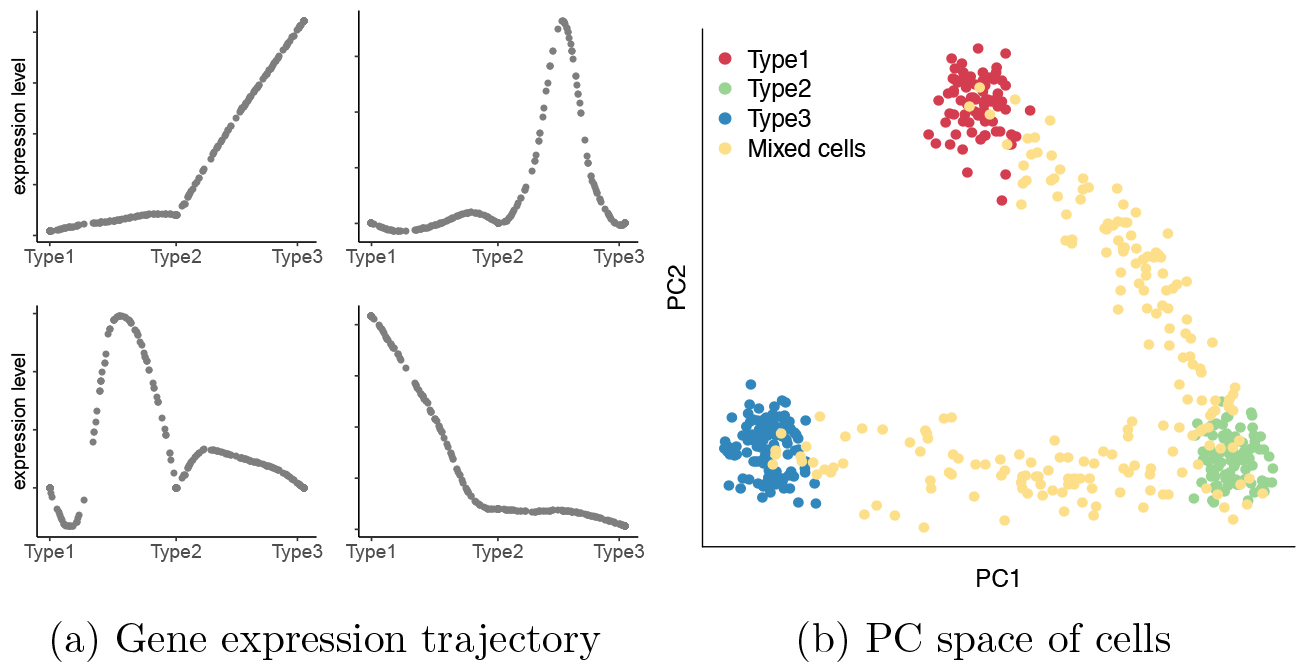
Illustration of the SOUP framework for 3 cell types with simulated developmental trajectory of type1 → type2 → type3. **(a)** Example of 4 differentially expressed genes along the developmental trajectory, with potentially nonlinear differentiation patterns. **(b)** Simulated 300 pure cells and 200 mixed cells, visualized in the leading principal component space.

Similar factorization problems as eq.(2) have appeared in previous literature under different settings. The most popular are the mixed-membership stochastic block model (MMSB) Mao et al. (2017) and topic modeling (for example, Arora et al. (2012, 2013); Huang et al. (2016)). However, it is nontrivial to extend these algorithms to our scenario. Similar formulation also appeared in Non-negative Matrix Factorization (NMF), where non-negative rank-*K* matrices Θ and *C* are estimated such that *X* ≈ Θ*C^T^*, for example, by minimizing the Euclidean distance (Lee & Seung, 2001). However, traditional NMF differs from our setting in two important ways: (i) the NMF problem is non-identifiable without introducing nontrivial assumptions, and (ii) SOUP does not rely on the non-negativeness of *C*, which makes it more broadly applicable to scRNA-seq data after certain preprocessing steps, such as batch-effect corrections, which can result in negative values. Recent work in Bing et al. (2017) considered the problem of overlapping *variable* clustering under latent factor models. Despite the different setup, the model comes down to a problem similar to eq.(2), and the authors proposed the LOVE algorithm to recover the variable allocation matrix, which can be treated as a generalized membership matrix. LOVE consists of two steps: (i) finding pure variables, and (ii) estimating the allocations of the remaining overlapping variables. Both steps rely on a critical tuning parameter that corresponds to the noise level, which can be estimated using a cross validation procedure. When we applied the LOVE algorithm to our single cell datasets, however, we found it sensitive to noise, leading to poor performance (Supplementary Section S3.5). Nonetheless, inspired by the LOVE algorithm, SOUP works in a similar two-step manner, while adopting different approaches in both parts. Most importantly, SOUP parameters are intuitive to set, and it is illustrated to have robust performance in both simulations and real data.

#### SOUP Algorithm

The SOUP algorithm involves finding the set of pure cells, and then estimating Θ. Pure cells play a critical role in this problem. Intuitively, they provide valuable information from which to recover the cluster centers, which further guides the estimation of Θ for the mixed cells. In fact, it has been shown in Bing et al. (2017) that the existence of pure cells is essential for model (2) to be identifiable, and we restate the Theorem below.

##### Theorem 1 (Identifiability)

*Model (2) is identifiable up to the permutation of labels, if (a)* Θ *is a membership matrix; (b) there exist at least 2 pure cells per cluster; and (c) Z is full rank.*

These assumptions are minimal, because in most single cell datasets, it is natural to expect the existence of at least a few pure cells in each type, and *Z* usually has larger entries along the diagonal.

The details of SOUP are presented in Methods and Supplemental Information. As an overview, to recover the pure cells the key is to notice the special block structure formed by the pure cells in the similarity matrix *A*. SOUP exploits this structure to calculate a purity score for each cell. This calculation requires two tuning parameters: *ϵ*, the fraction of most similar neighbors to be examined for each cell, and *γ*, the fraction of cells declared as pure after ranking the purity scores. After selection, the pure cells are partitioned into *K* clusters, by standard clustering algorithms such as K-means. The choice of *K* is guided by empirical investigations, including a sample splitting procedure (Supplementary Section S2).

To recover Θ, consider the top *K* eigenvectors of the similarity matrix *A*, denoted as *V* ∈ ℝ^*n*×*K*^. There exists a matrix *Q*\* ∈ ℝ^*K×K*^, such that Θ = *V Q*\* . If we have identified the set of pure cells ℐ and their partitions {*ℐ_k_*}, we essentially know their memberships, Θ_*ℐ·*_. Then it is straightforward to recover the desired *Q*\* from the sub-matrix Θ_*ℐ·*_= *V_ℐ·_Q*\*, which further recovers the full membership matrix Θ = *V Q*\* (Theorem 2). In practice, we plug in the sample similarity matrix 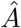 to obtain an estimate 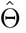, and we can further estimate 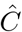 by minimizing 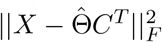.

##### Theorem 2 (SOUP clustering)

*In model (2), let V* ∈ ℝ*^n×K^ be the top K eigenvectors of A, and ℐ be the set of pure cells. Under the same assumptions as Theorem 1, the optimization problem*

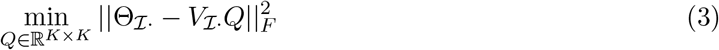

*has a unique solution Q* such that* Θ = *V Q\**.

The majority membership probability is max_*j*_ *θ_ij_*, and the majority type is the class that achieves the maximum.

#### Developmental Trajectories

SOUP provides two outcomes not available from hard clustering procedures such as (Kiselev et al., 2017; Lin et al., 2017; Satija et al., 2015): soft membership probabilities, 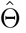, and soft cluster centers, 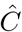. The next step is to estimate one or more developmental trajectories from the cells. Various algorithms have been developed that can identify multi-branching developmental trajectories in single cell data (Bendall et al., 2014; Shin et al., 2015; Ji & Ji, 2016; Setty et al., 2016; Street et al., 2018), and one successful direction is to estimate the lineages from cell clusters, usually by fitting a minimum spanning tree (MST) to the cluster centers in a low-dimensional space (Shin et al., 2015; Ji & Ji, 2016; Street et al., 2018), and then fitting a smooth branching curve to the inferred lineages (Street et al., 2018). It is straightforward to extend this idea to SOUP, where we identify the MST using SOUP estimated soft cluster centers, 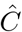. Following the common practice, 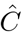 can be projected to a low-dimensional space for MST estimation. Notably, soft clusters provide an alternative input for Slingshot (Street et al., 2018), which yields more refined insights into development by providing less biased estimates of cluster centers in developing cells.

### Performance Evaluation

#### Simulations

There are no direct competitors of SOUP for semi-soft clustering in the single-cell literature, and here we use the following three candidates for comparison:

- Non-negative Matrix Factorization (NMF), where we use the standard algorithm from Lee & Seung (2001) to solve for non-negative 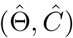 by

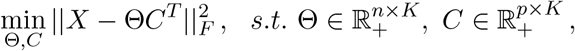

and we further normalize the solution 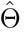 to be a proper membership matrix with unit row sums.
- Fuzzy C-Means (FC) (Bezdek, 1981), a generic soft clustering algorithm. Its tuning parameter *m* > 1 controls the cluster fuzziness, where *m* = 1 gives hard clustering. Here we present the results of the default choice *m* = 2.
- DIMMSC (Sun et al., 2017), a probabilistic clustering algorithm for single cell data based on Dirichlet mixture models. It is designed for hard clustering, but internally estimates the posterior probability of each cell belonging to different clusters, which can be treated as an estimator of Θ.

Although SOUP is derived from a linear model, it is robust and applicable to general scRNA-seq data. To illustrate this, we use the splat algorithm in the Splatter R package (Zappia et al., 2017) to conduct simulations. Splatter is a single-cell simulation framework that generates synthetic scRNA-seq data with hyper-parameters estimated from a real dataset. The algorithm incorporates expected violations of the model assumptions (see Methods). We simulate 500 genes and 300 pure cells from 4 clusters. Mixed cells are simulated along a developmental path and the number varies from 100 to 500. Throughout this section, we use the true *K* = 4 as input, and the default parameters for SOUP (*ϵ* = 0.1, *γ* = 0.5).

##### Soft membership estimation

For comparable evaluation across different scenarios with different cell numbers, we present the average *L*_1_ loss per cell, i.e., 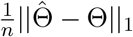, where ||·||_1_ is the usual vector *L*_1_ norm after vectorization. SOUP achieves the best performance under all scenarios (Figure 2a). In particular, with 100, 300, and 500 mixed cells, the true proportions of pure cells in the data are 75%, 50%, and 37.5%, respectively. Note that we always set *γ* = 0.5 for SOUP, which represents a prior guess of 50% pure cells, and we see that SOUP remains stable even when the given *γ* clearly overestimates or underestimates the pure proportion.

**Figure 2:**
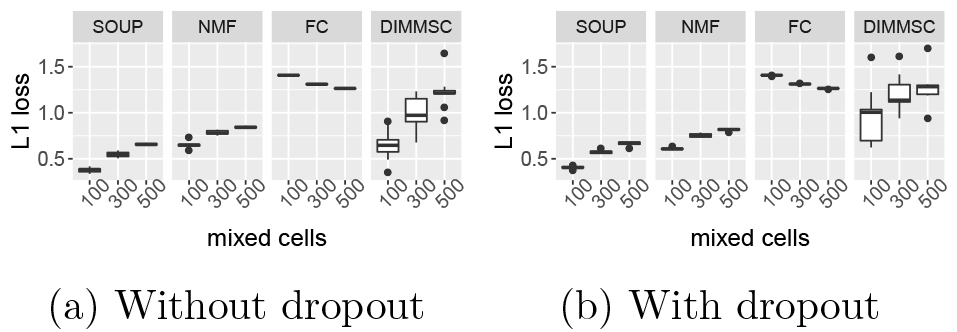
Boxplot of the average *L*_1_ losses of estimating Θ in 10 repetitions. Using the splat algorithm in the Splatter package, expression levels of 500 genes are simulated for 300 pure cells from 4 clusters, as well as {100, 300, 500} mixed cells along the trajectory of type1→type2→{type3 or type4}. **(a)** Without dropout; **(b)** with dropout.

##### Robustness to dropouts

One of the biggest challenges in single cell data is the existence of dropouts (Kolodziejczyk et al., 2015), where the mRNA for a gene fails to be amplified prior to sequencing, producing a “false” zero in the observed data. Here, we also evaluate the performance when the data is simulated with zero inflations, where the dropout parameters are also estimated from real data (see Splatter (Zappia et al., 2017) for details). We see that SOUP remains robust and outperforms all other algorithms (Figure 2b).

### SOUP as hard clustering

Although SOUP aims at recovering the full membership matrix Θ, it can also be used as a hard clustering method by labeling each cell as the majority type. In this final section, we benchmark SOUP as a hard clustering method on 7 labeled public single-cell datasets (Baron et al. (2016); Darmanis et al. (2015), details in Table S6). We compare SOUP to three popular single-cell clustering algorithms: (i) SC3 (Kiselev et al., 2017), (ii) CIDR (Lin et al., 2017), and (iii) Seurat (Satija et al., 2015). Because we aim at hard clustering, here we set *γ* = 0.8 for SOUP. We give the true *K* as input to SC3, CIDR, and SOUP. For Seurat, we follow the choices in Yang et al. (2017) and set the resolution parameter to be 0.9, and use the estimated number of principal components (nPC) from CIDR for Seurat. Even for hard clustering, SOUP is among the highest (Figure 3, showing Adjusted Rand Index (ARI)). Finally, we point out that when using the default choice of *γ* = 0.5, SOUP also achieves sensible performance, sometimes with even higher ARI (Table S6).

**Figure 3:**
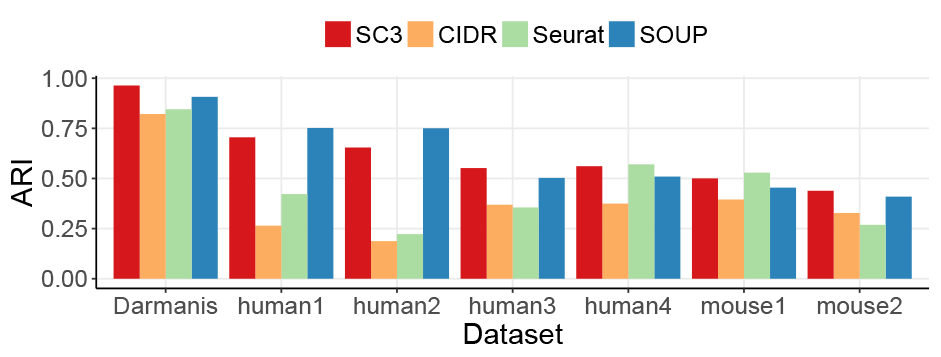
Adjusted Rand Index (ARI) on 7 labeled public datasets (Baron et al., 2016; Darmanis et al., 2015), using (i) SC3, (ii) CIDR, (iii) Seurat, and (iv) SOUP.

### Case Studies

#### Fetal brain cells I

We apply SOUP to a fetal brain scRNA-seq dataset, with 220 developing fetal brain cells between 12-13 gestational weeks (GW) (Camp et al., 2015). Guided with marker genes, these single cells are labeled with 7 types in the original paper: two subtypes of apical progenitors (AP1, AP2), two subtypes of basal progenitors (BP1, BP2), and three subtypes of neurons (N1, N2, N3). We refer to these as Camp labels. At this age many cells are still transitioning between different types, providing valuable information regarding brain development. Therefore, instead of the traditional hard clustering methods, SOUP can be used to recover the fine-scaled soft clustering structure.

We run SOUP with *K* = 2, 3, …, 7 on the log transformed transcript counts, and examine the clusters of cells, initially treating this as a hard clustering problem, and focusing on the dominating type for each cell. For *K* = 6 and 7, some clusters have no cells assigned to them, which is indicative of a misspecified *K*. For *K* = 5, the algorithm identifies cell types that correspond to A1, A2, B1, N2 and N3 in Camp’s nomenclature (Figure S6a, Figure 4 bottom right panel). For these data, when cells are in various developmental stages, hard clustering appears to overfit the data.

**Figure 4:**
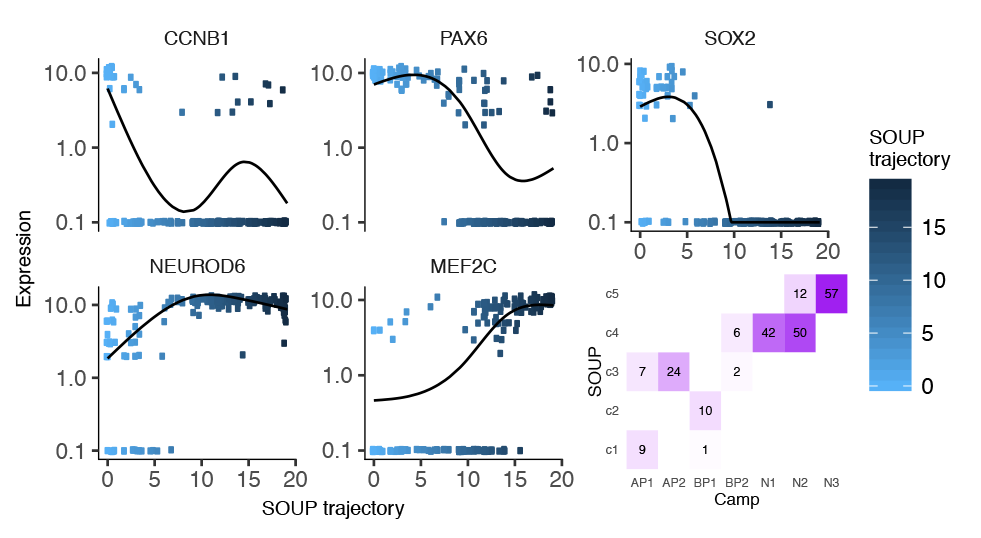
Expression levels of 5 anchor genes, visualized in log scale, where the 220 fetal brain cells are ordered by a SOUP uni-lineal developmental trajectory. The smooth lines are fitted by natural cubic splines with 3 degrees of freedom.

Next, we examine the soft assignments. For each cluster *k*, we label it by an anchor gene, which is the marker gene defined in Camp et al. (2015) that has the largest anchor score: [*C_gk_* − max{*C_g_*,_(−*k*)_}]/*sd*(*C_g_*,_(−*k*)_), where *C_g_*,_(−*k*)_ represents the center values of gene *g* on the (K-1) clusters other than *k*. The expression levels of the 5 anchor genes along the SOUP trajectory vary smoothly over developmental time (Figure 4), consistent with eq. (1). In the top 3 principal component space, the cells show a smooth developmental trajectory between clusters (Figure 5a), which is also consistent with eqs. (1-2).

**Figure 5:**
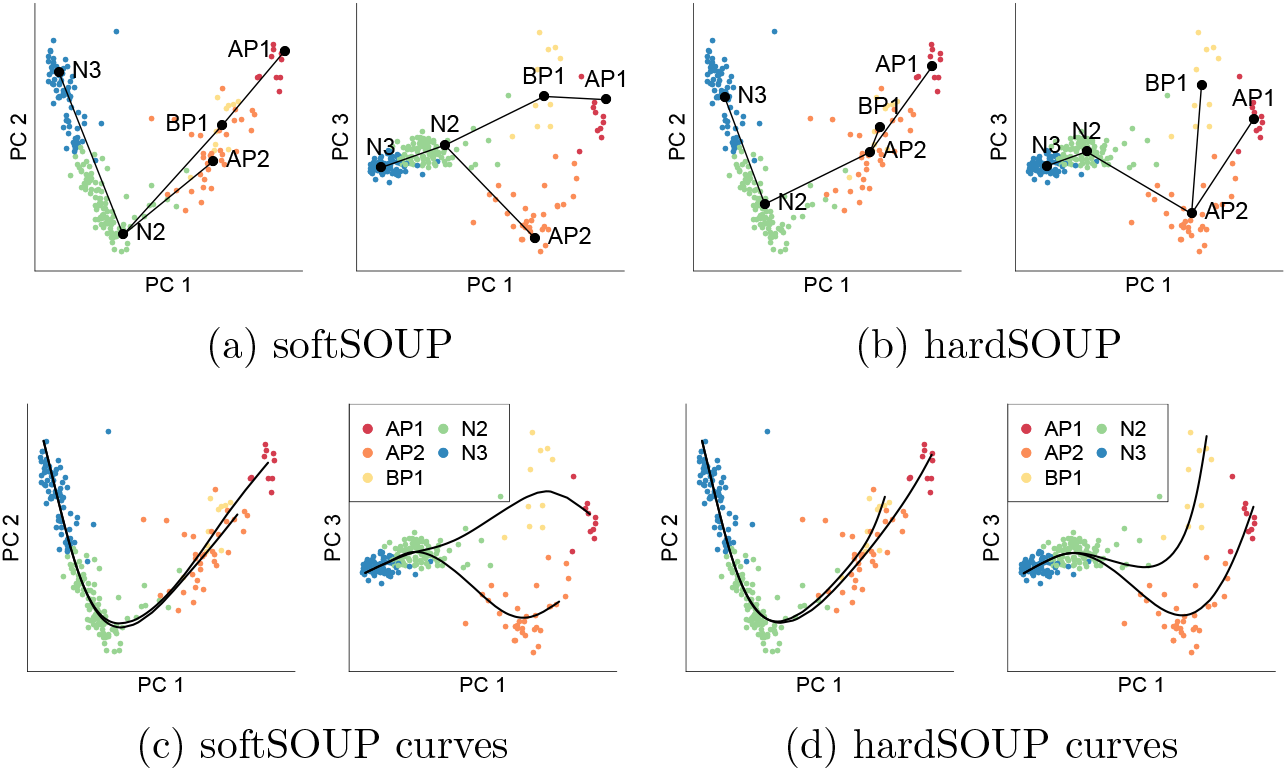
220 fetal brain cells, cluster centers, lineages, and branching curves in the top 3 principal component space. Cells are colored according to their SOUP major types, but annotated using Camp labels based on the largest overlap (see Figure 4). **(a,b)** MST of softSOUP and hard-SOUP cluster centers. **(c,d)** Smooth branching curves fitted by Slingshot based on MST in **(a,b)**, respectively.

To model the developmental trajectories we plot the cluster centers determined directly by SOUP (softSOUP) and by hard clustering (hardSOUP). Fitting a MST to the cluster centers, softSOUP identifies two lineages: AP-BP-N and AP-N (Figure 5a), both of which were previously described in Camp et al. (2015), while hardSOUP identifies less intuitive BP-AP-N and AP-N lineages (Figure 5b). Using Slingshot to fit smooth branching curves to these lineages via simultaneous principal curves, hardSOUP, recovers AP-N and BP-N transitions, and the artificial BP1-AP2 transition in the initial MST fit is dropped (Figure 5d). However, the AP-BP transition is still missing. softSOUP MST successfully reveals AP-N and AP-BP-N transitions (Figure 5a and 5c), thus capturing the true transition of cell types leading to neurons by accounting for the soft membership structures.

#### Fetal brain cells II

We next applied SOUP to a richer data set with 2,309 single cells from human embryonic prefrontal cortex (PFC) from 8 to 26 GW (Zhong et al., 2018). Using the Seurat package (Satija et al., 2015) the authors identified six major clusters: neural progenitor cells (NPC), excitatory neurons (EN), interneurons (IN), astrocytes (AST), oligodendrocyte progenitor cells (OPC) and microglia (MIC), which are referred to as Zhong labels. Our objective is to evaluate the developmental trajectories of the major cell types, after excluding IN and MIC, which are known to originate elsewhere and migrate to the PFC (Zhong et al., 2018). After several iterations of hard clustering by SOUP to remove IN and MIC cells (Tables S1-S3) 1503 cells remain, and they cluster into *K* = 7 types. These types correspond fairly well with the Zhong labels (Figure 6a); however, many cells have low majority membership probabilities (Figure S8) and do not strongly favor a particular cluster (Table S4). To illustrate this feature we display cells assigned to clusters 3 (NPC) and 7 (EN), color coded by the majority membership probability (Figure 6b). The two clusters divide the PC space evenly, with the pure cells identifying the cluster centers, while many non-pure cells can be best described as transitioning between clusters. SOUP captures the transitional nature by soft clustering.

**Figure 6:**
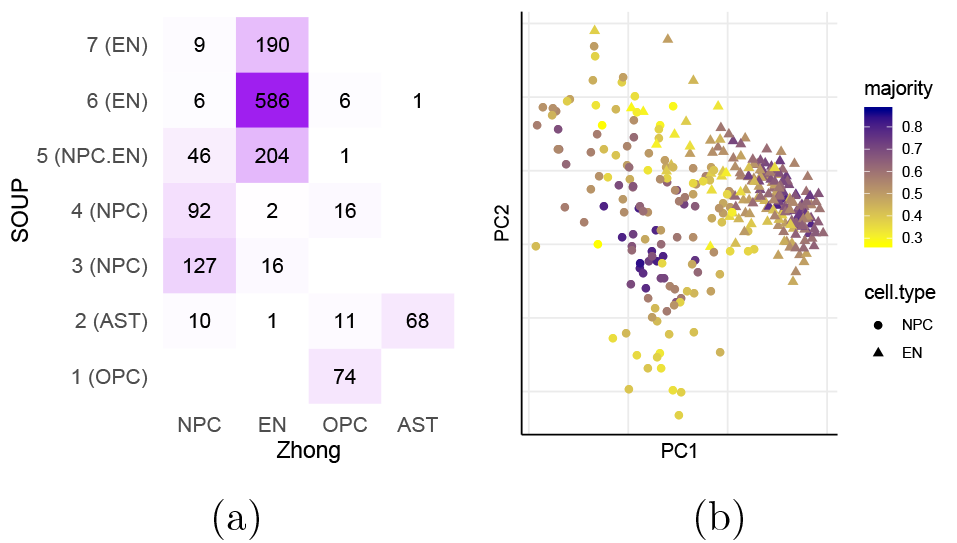
**(a)** Contingency table of Zhong labels and major SOUP labels excluding IN and MIC. (b) Distribution of cluster 7 (EN) and cluster 3 (NPC) cells and their majority membership probabilities.

The SOUP trajectories reveal two developmental paths (Figure 7): a neuronal lineage showing NPCs evolving to ENs (clusters: 4 → 3 → 7 → 6 → 5) and a glial lineage showing NPCs evolving to OPCs and then to ASTs. Projecting the cells onto the lineages can provide pseudotime estimates of development. The lineages correspond roughly with sampled GWs (Table S4). Our results are similar to those in (Zhong et al., 2018), however we found that NPCs evolve to OPCs and then to ASTs (clusters: 4 → 3 → 1 → 2). The latter transitional step, which differs from the published analysis, is consistent with the literature (Zhu et al., 2008). Finally, cluster 5, which consists of a mixture of cells Zhong labeled as EN and NPC, is placed at the end of the neuronal lineage, suggesting that some of the NPC labels are incorrect and that this cluster constitutes a distinct class of ENs.

**Figure 7:**
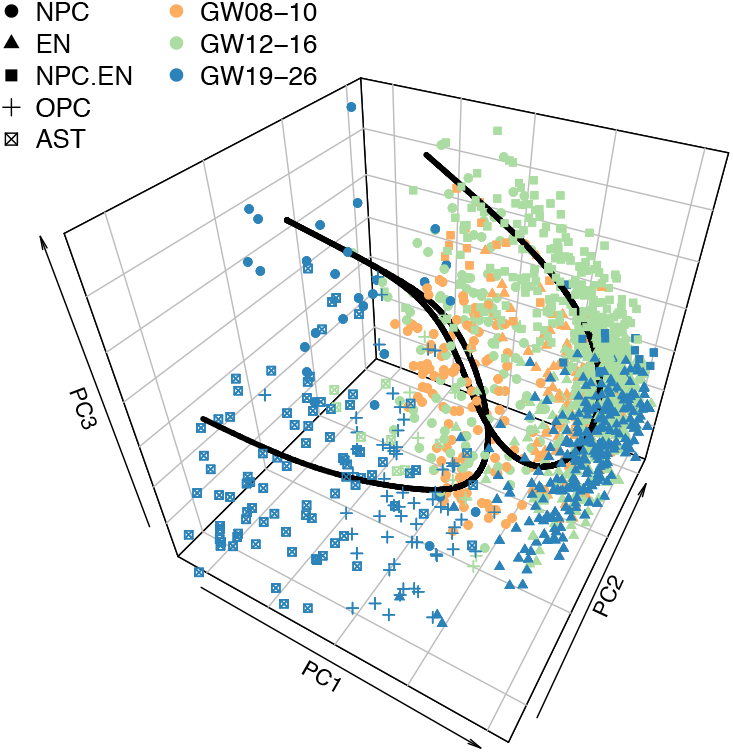
Developmental trajectories of 1503 Zhong cells delineate glial and neuronal pathways. Cluster labels are defined in Figure 6a.

Additional strengths of SOUP are highlighted by analyses described in Supplement, which investigate gene expression as a function of cell membership to cluster and the proximity of cells to the neuronal trajectory (Figures 7,S9). In particular we evaluate the final clusters of the neuronal lineage, clusters 5 and 6. In terms of gene expression, cells in cluster 6 shows all the hallmarks of neuronal development, including low expression of neuronal markers in immature and much higher expression in maturing neurons. There is also some evidence of heterogeneity of expression of genes marking neurons in some cells, consistent with differentiation into different neuronal subtypes. For cells from cluster 5, the evidence is far less clear: the majority of cells manifest neuronal markers at high levels, consistent with maturing neurons; yet, there is also expression of substantial set of NPC markers in these neurons, a puzzling feature that could be either a technical artifact or an unanticipated developmental feature of deep layer projection neurons.

## Discussion

We develop SOUP, a novel semi-soft clustering algorithm for single cell data. SOUP fills the gap of modeling uncertain cell labels, including cells that are transitioning between cell types, which is ubiquitous in single cell datasets. SOUP outperforms generic soft clustering algorithms and, if treated as hard clustering, it also achieves comparable performance as state-of-the-art single cell clustering methods. By using soft clustering input, it can provide an estimate of developmental trajectories that is less biased and these results reflect valuable information regarding developmental patterns. We present the results from two case studies based on expression of human fetal brain cells and find SOUP reveals patterns of development not apparent in prior published analyses.

As is typical for clustering algorithms, selecting the optimal number of clusters, *K*, is challenging. We recommend balancing input from several empirical approaches and iterating over a range of *K* to determine a good choice. Notably, applying SOUP to a different numbers of clusters reveals hierarchical structure among the cell types. To determine fine scale structure within major cell types, SOUP can be applied iteratively to subsets of cells.

Using SOUP to obtain soft membership probabilities and then estimating developmental trajectories provides two complementary views of the data. Some cells can be reliably assigned to a cluster and these cells constitute pure types, which can be highly informative. Other cells are transitioning and estimated membership will fall within two, or even more cell types. Examining the membership probabilities, and the placement on a developmental trajectory, provides critical information about the developmental processes and offers a parsimonious and scientifically meaningful alternative to estimating a large number of discrete cell types.

Notably, although SOUP is derived under a generic additive noise model and does not explicitly model the technical noise such as dropouts, we find it to be robust when applied to realistic simulations and to a variety of single cell datasets. Moreover, it is computationally efficient, which makes it easily applicable to large single cell datasets. SOUP takes less than 15 minutes for 3,600 cells and 20,000 genes, benchmarked on a linux computer equipped with AMD Opteron(tm) Processor 6320 @ 2.8 GHz. Therefore, SOUP is a versatile tool for single cell analyses.

## Methods

### SOUP

Our SOUP algorithm contains two steps: (i) find the set of pure cells, and (ii) estimate Θ. Pure cells play a critical role in this problem. Intuitively, they provide valuable information from which to recover the cluster centers, which further guides the estimation of Θ for the mixed cells. Once the pure cells are identified then the algorithm proceeds as described in Results.

#### Find pure cells

Denote the set of pure cells in cluster *k* as

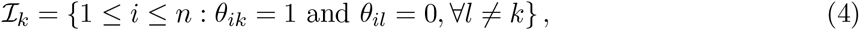

and the set of all pure cells as 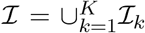. To recover ℐ, the key is to notice the special block structure formed by the pure cells in the similarity *A*. In particular, under eq.(2), the pure cells form *K* blocks in *A*, where the entries in these blocks are also the maxima in their rows and columns, ignoring the diagonal. Specifically, define

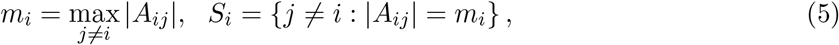

and we call *S_i_* the *extreme neighbors* of cell *i*. It can be shown that if cell *i* is pure, then |*A_ij_*| = *m_i_* = *m_j_* for all *j* ∈ *S_i_*. On the contrary, for a mixed cell *i*, there exist some cells *j* ∈ *S_i_* where *m_j_* > |*A_ij_*|. Inspired by these observations, we define a purity score of each cell,

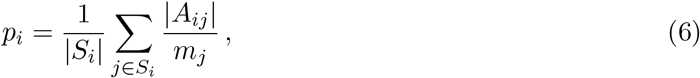

then naturally *p_i_* ∈ [0, 1]. Furthermore, the pure cells have the highest purity scores, that is, ℐ = {*i*: *p_i_* = 1} (Theorem S1).

In practice, we plug in the sample similarity matrix 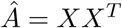, and estimate *S_i_* and *p_i_* by

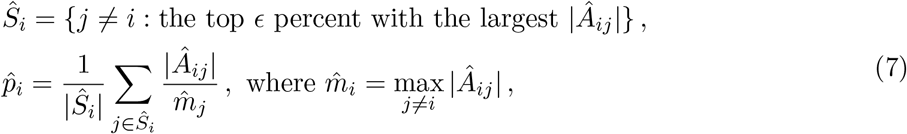

and we estimate ℐ with the top *γ* percent of cells:

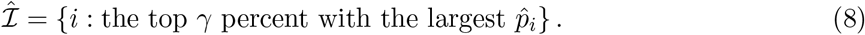

Finally, these pure cells are partitioned into *K* clusters, 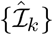, by standard clustering algorithms such as K-means. The complete algorithm is summarized in Supplementary S1.1.

#### Tuning parameters

The two tuning parameters of SOUP are the quantiles, *ϵ* and *γ*, both intuitive to set. The quantile *γ* should be an estimate of the proportion of pure cells in the data, of which we usually have prior knowledge. In practice, we find that SOUP remains stable even when *γ* is far from the true pure proportion, and it is helpful to use a generous choice. Throughout this paper, we always set *γ* = 0.5 and obtain sensible results. As for *ϵ*, it corresponds to the smallest proportion of per-type pure cells, and it suffices if *ϵ* ≤ min_*k*_ |*ℐ_k_*|/*n*, so that 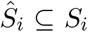 for pure cells. This choice does not need to be exact, as long as *ϵ* is a reasonable lower bound. In practice, we find it often beneficial to use a smaller *ϵ* that corresponds to less than 100 pure cells per type. By default, we use *ϵ* = 0.1 for datasets with less than 1, 000 cells, *ϵ* = 0.05 for 1,000 - 2,000 cells, and *ϵ* = 0.03 for even larger datasets. Simulation results of sensitivity are presented in Supplementary Section S3.4.

#### Gene selection

It is usually expected that not all genes are informative for clustering. For example, housekeeping genes are unlikely to differ across cell types, hence provide limited information for clustering other than introducing extra noise. Therefore, it is desirable to select a set of informative genes before applying SOUP clustering. Here, we combine two approaches for gene selection: (i) the DESCEND algorithm proposed in Wang et al. (2017) based on the Gini index, and (ii) the Sparse PCA (SPCA) algorithm (Witten et al., 2009) (see Supplementary Section S1.2 for details).

### Simulations

We conduct simulations using the splat algorithm in the Splatter R package (Zappia et al., 2017). Splatter estimates the simulation parameters from a real dataset, and can generate synthetic scRNA-seq data with cells from multiple populations or along differentiation paths. Here, we use the Zeisel data (Zeisel et al., 2015) for hyper-parameter estimation, where splat has shown to be successful (Zappia et al., 2017).

We simulate 500 genes and 300 pure cells from 4 clusters with probability (0.2, 0.2, 0.2, 0.4). By default, Splatter randomly selects 10% of the genes to have differential expression levels among cell types, and the separation is controlled by the location factor, `deFactor`, where larger value leads to more distinct cell types. Here, we present the results when `deFactor`=2, and more scenarios can be found in the Supplement.

We simulate different numbers of mixed cells using the “path” method in Splatter. In particular, we consider the trajectory structure where cell type 1 differentiates into an intermediate cell type 2, which then differentiates into two mature cell types, 3 and 4. Each mixed cell is randomly placed along one of the three paths with probability (0.3, 0.3, 0.4). Each path is a continuous development from a starting cluster to an ending cluster, and the expression level of each gene changes either linearly or nonlinearly from the starting expression level to the ending expression level (examples in Figure 1a). Here, we use the default setting where 10% of the genes have nonlinear differentiation paths. More scenarios are presented in the Supplement.

All algorithms are applied to the log-transformed data, except for DIMMSC which is developed under a Multinomial model for count data. NMF can be applied to the raw count data as well, which usually has slightly worse performance. We also tried other choices of *m*’s for fuzzy C-Means, and obtained similar results. The complete comparison can be found in the Supplement.

## Acknowledgement

This work was supported by NIMH grants R37MH057881 and R01MH109900 and the Simons Foundation SFARI 402281 and 367561. The R package of SOUP is available at https://github.com/lingxuez/SOUPR.

## Supporting Information

### S1 Details of SOUP

#### S1.1 Algorithms

For the ease of readability, we summarize the two steps of SOUP in Algorithm 1 and Algorithm 2. Note that when solving for 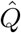 in Algorithm 2, theoretically one would consider the constrained problem:

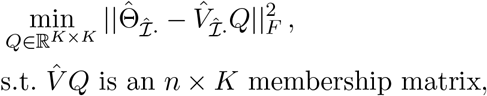

where membership matrices are the ones with nonnegative entries and unit row sums. However, in practice, solving this constrained problem is computationally demanding, sometimes with empty feasible sets. Therefore, we adopt a heuristic approach that first solves the unconstrained optimization problem as in Algorithm 2, and normalizes the obtained 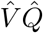 afterwards to get a membership matrix.

##### Algorithm 1 findPure

**Input:** similarity matrix 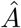, quantile *ϵ*, quantile *γ*

**Output:** estimated set of pure cells 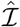

1: For each cell *i*,

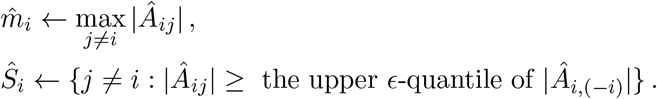
2: For each cell *i*, compute its purity score:

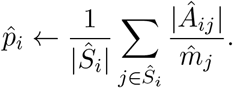
3: 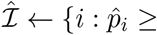 the upper *γ*-quantile of 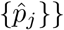.

#### S1.2 Gene selection

It is usually expected that not all genes are informative for clustering. For example, housekeeping genes are unlikely to differ across cell types, hence provide limited information for clustering other than noise. Therefore, it is desirable to select a set of informative (i.e. highly variable) genes before applying SOUP. Here, we combine two selection approaches. The first is the DESCEND method proposed in Wang et al. (2017), where the authors developed a semi-parametric approach to estimate several distribution statistics for each gene, including the Gini index that indicates the excessive variability of gene expression levels across cells. The authors suggested to threshold the normalized difference between the observed and expected Gini index, and use the set of highly variable genes for clustering. Throughout this paper, we use the default threshold of 3. We refer the readers to the original paper (Wang et al., 2017) for more details.

##### Algorithm 2 SOUP clustering

**Input:** data *X*, number of clusters *K*, quantile *ϵ*, quantile *γ*

**Output:** estimated membership 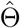

1: 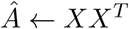
2: 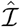 ← *findPure*(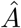, *K*, *ϵ*, *γ*), and apply K-means clustering on 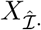 to get the partition 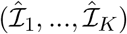.
3: 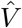 ← the top *K* eigenvectors of 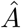.
4: Let 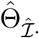 be the membership submatrix for pure cells, obtained by putting 1’s and 0’s in the corresponding columns according to 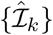. Solve for

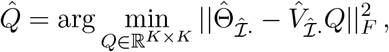

where ||·||*_F_* is the Frobenius norm.
5: 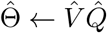.

The second approach is based on Sparse Principal Component Analysis (SPCA). In fact, it is common to first apply PCA, and select genes with the highest loadings in the top few principal components (PCs) for clustering (see Camp et al. (2015) for an example). SPCA provides a more rigorous algorithm to implement this idea, where it directly solves for the leading *sparse* PCs, and genes with nonzero entries are selected. In this paper, we use the efficient SPCA algorithm in Witten et al. (2009). The algorithm requires a tuning parameter, *c*, that controls the sparsity of the solution, where smaller *c* leads to sparser results, hence fewer selected genes. Throughout this paper, we always set *c* = 0.05, and use the top three sparse PCs.

In our experiments, we find that DESCEND and SPCA usually capture different structures in the data. SPCA usually picks up genes that differentiate major cell types, while DESCEND usually identifies genes that distinguish finer scaled clustering structures. Therefore, the best performance is achieved by combining both lists of genes.

#### S1.3 Normalization

In practice, cells can have different scaling due to sequencing depths and cell sizes, and proper normalization is required prior to using SOUP. Formally, we have 𝔼[*X_raw_*] = Diag((*s_i_*))Θ*C^T^*, where *s_i_* is the scaling factor of cell *i*. The factor *s_i_* can be interpreted as efficiency, while *C* is the expected expression level of each type. Alternatively, *s_i_* can represent the library size, with *C* being the *relative* type-specific expression profile. This factor can be estimated in various ways, sometimes with the help of spike-ins (Wang et al., 2017). Here, for RNA-seq data, we simply treat *s_i_* as the library size, and normalize such that the total sum of counts, ∑_*g*_ X_*ig*_, is 10^6^ in each cell *i*, which is essentially the Transcript per Million (TPM) normalization. In practice, we find it usually beneficial to further apply a log-transformation before running SOUP.

#### S1.4 SOUP for count data

The SOUP algorithm is derived under a generic additive noise model (eq.(1)). Here, we point out that SOUP is applicable to more general scenarios as long as we construct a similarity matrix that has structures similar to eq.(2). In particular, under a Poisson model, we have

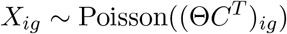

for cell *i* and gene *g* independently, we can compute the following similarity matrix

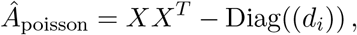

where *d_i_* = ∑_*g*_ *X_ig_*. Then it can be shown that

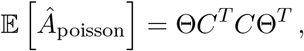

which has the same structure as eq.(2) with *σ*^2^ = 0, hence SOUP still applies.

However, when applied to single cell datasets, we find it usually beneficial to log-transform the RNA-seq data and use the general algorithm for additive noise model (Algorithm 1 and Algorithm 2). In addition, centering data with respect to genes when computing *A* is also helpful for identifying pure cells, because this usually makes *Z* more diagonally dominant, leading to a larger separation in purity scores between pure cells and mixed cells, as indicated in Theorem S1.

#### S1.5 Theorems and proofs

##### Theorem S1 (Pure cells)

*In model (2), under the same assumptions as Theorem 1, if we further require*

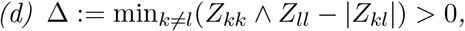

*then we have* ℐ = {*i*: *p_i_* = 1}*, where p_i_ is the purity score as defined in eq.(6).*

*Proof of Theorem S1.* Note that

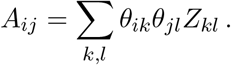

For any cell *i*, consider the two cases:

- If cell *i* is pure for type *k*, then for any *j* ≠ *i*,

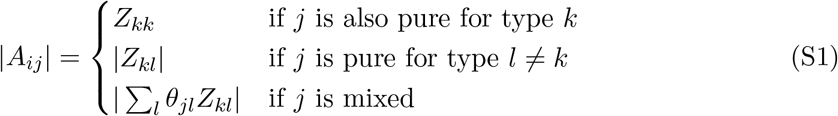 Note that |*Z_kl_*| ≤ *Z_kk_* − Δ, and we can also show that in the last case,

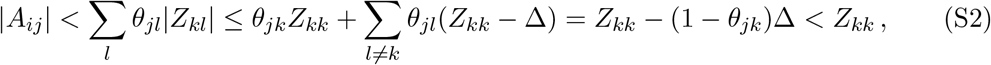

hence

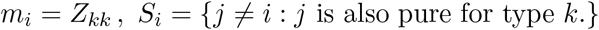
- If cell *i* is mixed, then for any *j* ≠ *i*,

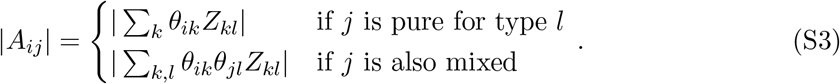 For the first case, we already shown above that

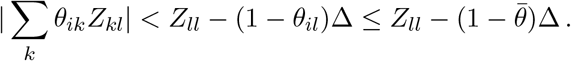 For the second case, we have

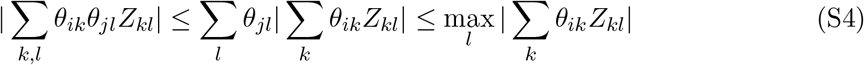 Therefore, *m_i_* is achieved at (*i, j*) for the pure cells *j* in some type *l*\*:

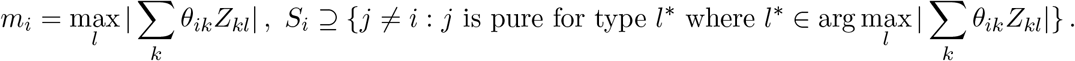

Therefore, the conclusions follow because

- if *i* is pure for type *k*, then

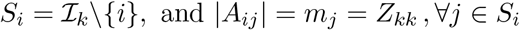

hence *p_i_* = 1.
- if *i* is mixed, then *S_i_* contains pure cells from cluster(s) *l*\* ∈ arg max_*l*_ |∑*_k_θ_ik_Z_kl_*|, and

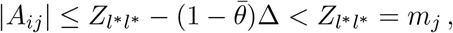

hence 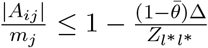 for all pure cells in *l\**, which further implies *p_i_* < *1.*

##### Proof of Theorem 1

See Theorem 2 in Bing et al. (2017). Note that Assumption (b), the existence of pure cells, is necessary for identifiability. We refer the readers to Bing et al. (2017) for an example of an unidentifiable model where assumptions (a) and (c) hold, but no pure cells exist.

##### Proof of Theorem 2

First, consider the noise free scenario, 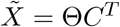, then we have

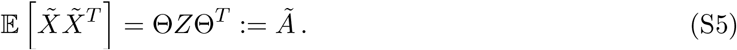

Consider the following symmetric decomposition problem,

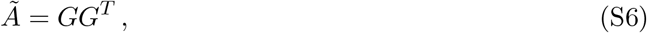

then Θ*Z*^1*/*2^ is one solution. Although the solution is not unique when *Z* is not diagonal, it has been shown in Mao et al. (2017) that under the same assumptions as in Theorem 1, for any solution *G*, there exists a *K* × *K* matrix *O*, such that

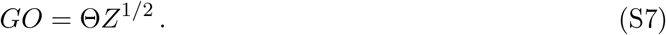

In particular, let 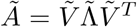 be the eigen decomposition of 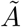, then 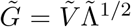 is one solution to problem (S6), which implies

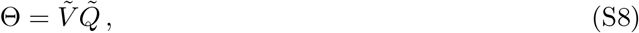

where 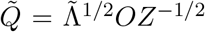 is a *K* × *K* matrix. In order to find 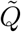, recall that we are given the set of pure cells ℐ and their partitions {ℐ*_k_*}. Equivalently, we already know the corresponding rows of the membership matrix, Θ_*ℐ·*_. Therefore, the desired 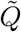 can be solved from

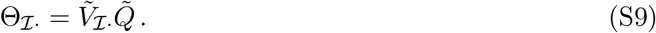

In general, under model (2), we have

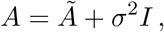

where 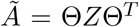 is the above noise-free similarity matrix. Let *V* be the top *K* eigenvectors for *A*, then 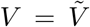, hence the above arguments also hold for *V* . Finally, the solution to problem (3) is unique because its Hessian matrix, (*V_ℐ·_*)^*T*^ *V_ℐ·_*, is positive definite due to assumption (b) in Theorem 1. Furthermore, the minimum is 0, achieved at 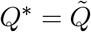, and Θ = *V Q*\*.

## S2 Selecting *K*

### S2.1 Selecting *K* from pure cells

One of the main challenges of SOUP is how to select the number of clusters, *K*. A simple solution is to focus only on the pure cells, treat it as a hard clustering problem, and apply standard selection techniques. For example, one can apply K-Means to the pure cells with a sequence of different *K*’s, and choose the optimal *K* according to certain metrics. Here, we examine the performance of two widely used metrics: (i) Calinski-Harabasz (CH) index (Caliński & Harabasz, 1974), which is used to identify the number of clusters in CIDR (Lin et al., 2017), and (ii) Silhoutte score (Rousseeuw, 1987), another popular metric. Both metrics select *K* that achieves the highest score, where CH index measures the normalized ratio between between-cluster and within-cluster variations, and Silhouette score measures how points are similar to its own cluster compared to other clusters. However, when applied to the single cell datasets, both metrics reveal only the major clusters, leading to *K* = 2 in the Camp dataset (Figure S1a and S1b). We will show that this is true in other public single-cell datasets as well (Table S7).

### S2.2 Selecting *K* with sample splitting

Sample splitting has been used to select the optimal rank for matrix completion (Kanagal & Sindhwani, 2010; Owen & Perry, 2009), and recently, similar idea has been extended to hard clustering for selecting the number of clusters (Fu & Perry, 2017). Here, we follow the bi-cross-validation procedure in Owen & Perry (2009); Fu & Perry (2017), and extend it to our soft clustering problem. Specifically, we permute the rows and columns of the expression matrix *X*, and partition them as *n* = *n*_1_ + *n*_2_ and *p* = *p*_1_ + *p*_2_. For the ease of presentation, assume *X* has been permuted, with partition

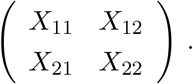

**Figure S1:**
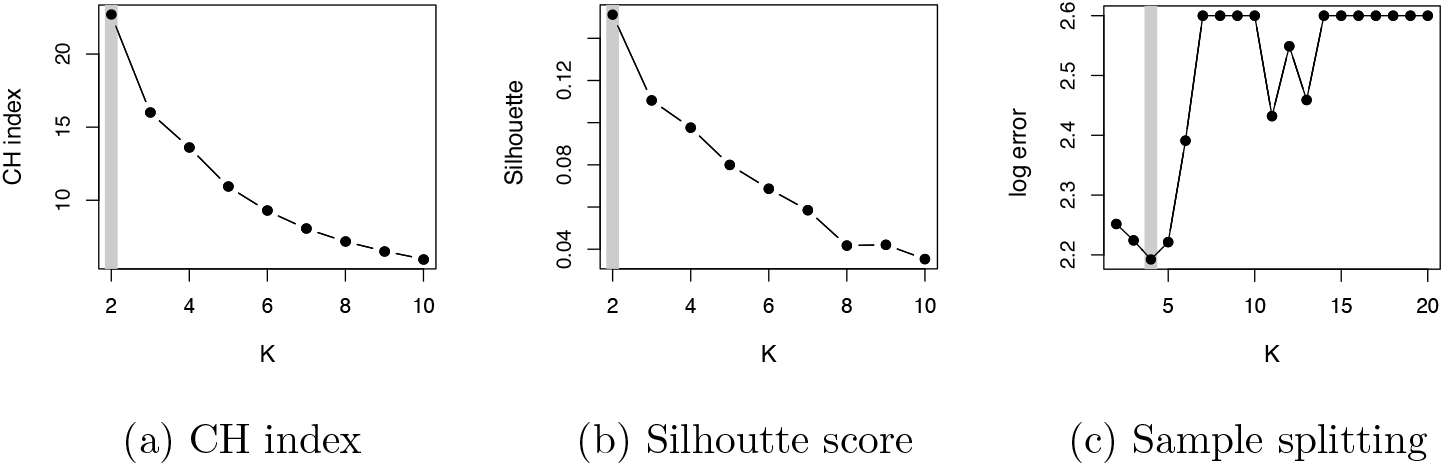
**(a)** Calinski-Harabasz (CH) index, **(b)** Silhoutte score, and **(c)** sample splitting prediction error for selecting the number of clusters in the Camp fetal brain data. CH index and Silhoutte score are computed using K-means hard-clustering results, applied on the pure cells identified by SOUP, and 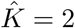 is selected. Sample splitting prediction errors are averaged across 10 repetitions. For visualization purpose, the values are plotted in the log scale and capped at 2.6.

We treat (*X*_21_*, X*_22_) ∈ ℝ^*n*_2_*×p*^ as the training samples, and *X*_11_ ∈ ℝ^*n*_1_*×p*_1_^ as the held out block. For each *k*, we compute the prediction error of *X*_11_ as below:

1. Apply SOUP to the training samples, (*X*_21_*, X*_22_), with *k* clusters, to get the estimated membership 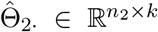 and center matrix 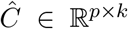. We partition 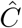 accordingly to get 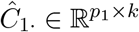 and 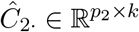.
2. Estimate the membership of the held out samples using *X*_12_ and 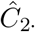. Specifically, we first solve for

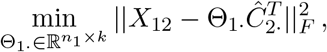

and then normalize to get the proper membership matrix 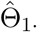.
3. Finally, the prediction of the held out block, *X*_11_, is obtained by 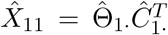, and the prediction error is computed as 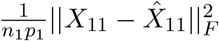.

In practice, we always split the rows and columns into equally sized partitions. This procedure is repeated a few times, and the *K* that achieves the smallest average prediction error is selected. We apply this procedure to the Camp data to search over *K* ∈ {2, …, 20}, where the prediction error is averaged over 10 repetitions, and obtain 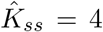 (Figure S1c). Note that *K* = 4 successfully distinguishes two subtypes of neuronal progenitors as well as early and matured neurons (Figure S6a).

As a final check on our choice of *K*, we examine the membership matrix *θ*. We require that at least 2 cells have majority probability > .5, for every cell type.

### S2.3 Benchmarking on public datasets

Finally, we benchmark different selection procedures in the 7 public datasets with known labels that were used in Section, and we focus on the set of selected informative genes via DESCEND and SPCA (Table S6). As in the main text, we use *γ* = 0.8 for SOUP to pick 80% of the cells as pure. We compare the selected *K* among {2, 3, …, 20} using (i) optimal CH index or Silhouette score computed with K-means hard-clustering results of the pure cells, and (ii) minimal prediction error in sample splitting. We then apply SOUP clustering using the selected *K*, and compute the Adjusted Rand Index (ARI), using the reference labels as gold standard (Table S7). Again, we see that standard selection metrics substantially underestimates the number of clusters, revealing only major split of cell types with poor ARI. On the contrary, the sample splitting procedure gives more sensible estimates that are close to the reference, and lead to similar ARI as if the true *K* is given. In fact, in the six pancreatic datasets from Baron et al. (2016), using 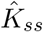 sometimes leads to even higher ARI than using the reference *K* (Table S6, Table S7). This is probably because in some datasets, certain reference cell types contain less than 5 cells, which are difficult to separate as distinct clusters.

## S3 Supplementary Simulations

### S3.1 Simulation settings

First, we describe the detailed simulation settings. We compare SOUP with (i) Non-negative Matrix Factorization (NMF) (Lee & Seung, 2001), applied on either count-scale (NMF-ct) or log-scale (NMF-log), where the solution is normalized to have unit row sum; (ii) Fuzzy C-means (FC) (Bezdek, 1981) with three choices of *m*, *m* ∈ {1.5, 2, 5}; (iii) DIMMSC (Sun et al., 2017), which is based on a Multinomial count model. We use the true *K* = 4 as input for all algorithms, and the default parameters for SOUP (*ϵ* = 0.1, *γ* = 0.5). Throughout this section, we do not perform gene selection. We apply normalization and log-transformation for NMF-log, SOUP and FC, and give the raw counts to NMF-ct and DIMMSC.

We conduct simulations using the splat algorithm in the Splatter R package (Zappia et al., 2017), where the Zeisel data (Zeisel et al., 2015) is used for hyper-parameter estimation. We simulate 500 genes and 300 pure cells from 4 clusters with probability (0.2, 0.2, 0.2, 0.4). By default, Splatter randomly selects 10% of the genes to have differential expression levels among cell types, and the separation is controlled by the location factor, `deFactor`, where larger value leads to more distinct cell types. Here, we present the results of `deFactor` ∈ {1, 2, 5}.

To evaluate the performance under various proportions of mixed cells, we simulate {100, 300, 500} mixed cells using the “path” method in Splatter. In particular, we consider the trajectory structure where cell type 1 differentiates into an intermediate cell type 2, which then differentiates into two mature cell types, 3 and 4. Each mixed cell is randomly placed along one of the three paths with probability (0.3, 0.3, 0.4). Each path is a continuous development from a starting cluster to an ending cluster, and the expression level of each gene can change either linearly or nonlinearly from the starting expression level to the ending level. By default, 10% of the genes have nonlinear differentiation patterns.

### S3.2 Soft membership estimation

We conduct 10 repetitions in each setting, and compare the *L*_1_ losses of estimating Θ. To achieve comparable evaluation across different scenarios with different cell numbers, we present the average loss per cell, i.e., 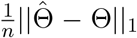, where ||·||_1_ is the usual vector *L*_1_ norm after vectorization. In Figure S2, we see that SOUP achieves the best performance regardless of the number of mixed cells and the cluster separation `deFactor`. In addition, note that we always set *γ* = 0.5 for SOUP, which represents a prior guess of 50% pure cells, but with *n_mix_* ∈ {100, 300, 500}, the true pure proportions are {0.75, 0.5, 0.375}, respectively. We see that SOUP is stable even when *γ* underestimates or overestimates the pure proportion.

We also evaluate the robustness of different algorithms to dropouts using the zero-inflated simulator of Splatter. The dropout parameters are also estimated from the Zeisel data. Again, we evaluate the average *L*_1_ loss of estimating Θ, and for simplicity, we present only the results with 500 mixed cells (Figure S2). We see that SOUP achieves robust performance with the existence of dropouts, and always outperforms other algorithms.

### S3.3 Robustness to nonlinear trajectories

Although SOUP is derived from a linear model, it is still applicable to general scenarios where genes exhibit nonlinear differentiation patterns along developmental trajectories. To illustrate this point, we use the Splatter package to simulate development paths where {10%, 30%, 50%, 70%} of the genes follow nonlinear development patterns from the starting cluster to the ending cluster. We refer the readers to the original paper (Zappia et al., 2017) of the detailed procedure of path simulation. The remaining settings are the same as before. Again, we evaluate the average *L*_1_ loss of SOUP when {100, 300, 500} mixed cells are simulated, with `deFactor`=2, with and without dropout. We see that SOUP is robust to such nonlinearity (Figure S3).

### S3.4 Sensitivity to tuning parameters

Here, we examine the sensitivity of SOUP to its two tuning parameters, *ϵ* and *γ*. Recall that *ϵ* represents the smallest proportion of per-type pure cells, min_*k*_ |*ℐ_k_*|/*n*, and *γ* represents the proportion of pure cells, |*ℐ*|/*n*. Following the previous simulation settings, we examine the performance of SOUP with the existence of dropouts and `deFactor`=5. Note that there are 300 pure cells, therefore, with {100, 300, 500} mixed cells, the optimal tuning parameters are *ϵ*\* ∈ {0.25, 0.167, 0.125} and *γ*\* ∈ {0.75, 0.5, 0.375}, respectively. We present the average *L*_1_ losses of estimating Θ when using *ϵ* ∈ {0.05, 0.1, 0.15, 0.2} and *γ* ∈ {0.3, 0.5, 0.7} (Figure S4). We see that SOUP is robust as long as *ϵ* < *ϵ*\* and *γ* ≈ *γ*\* . In particular, it is more important to choose a smaller *c* so that the estimated extreme neighbors, 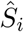, has a high precision in recovering *S_i_*. Therefore, by default, we use *ϵ*= 0.1 for datasets with less than 1,000 cells, *ϵ* = 0.05 for 1,000 - 2,000 cells, and *c* = 0.03 for even larger datasets. On the other hand, the performance is more robust to the choice of *γ*, where the default choice *γ* = 0.5 is usually sensible . In fact, it is usually beneficial to use a slightly more generous *γ*, and under this case, the performance is robust even when *ϵ* > *ϵ*\*.

**Figure S2:**
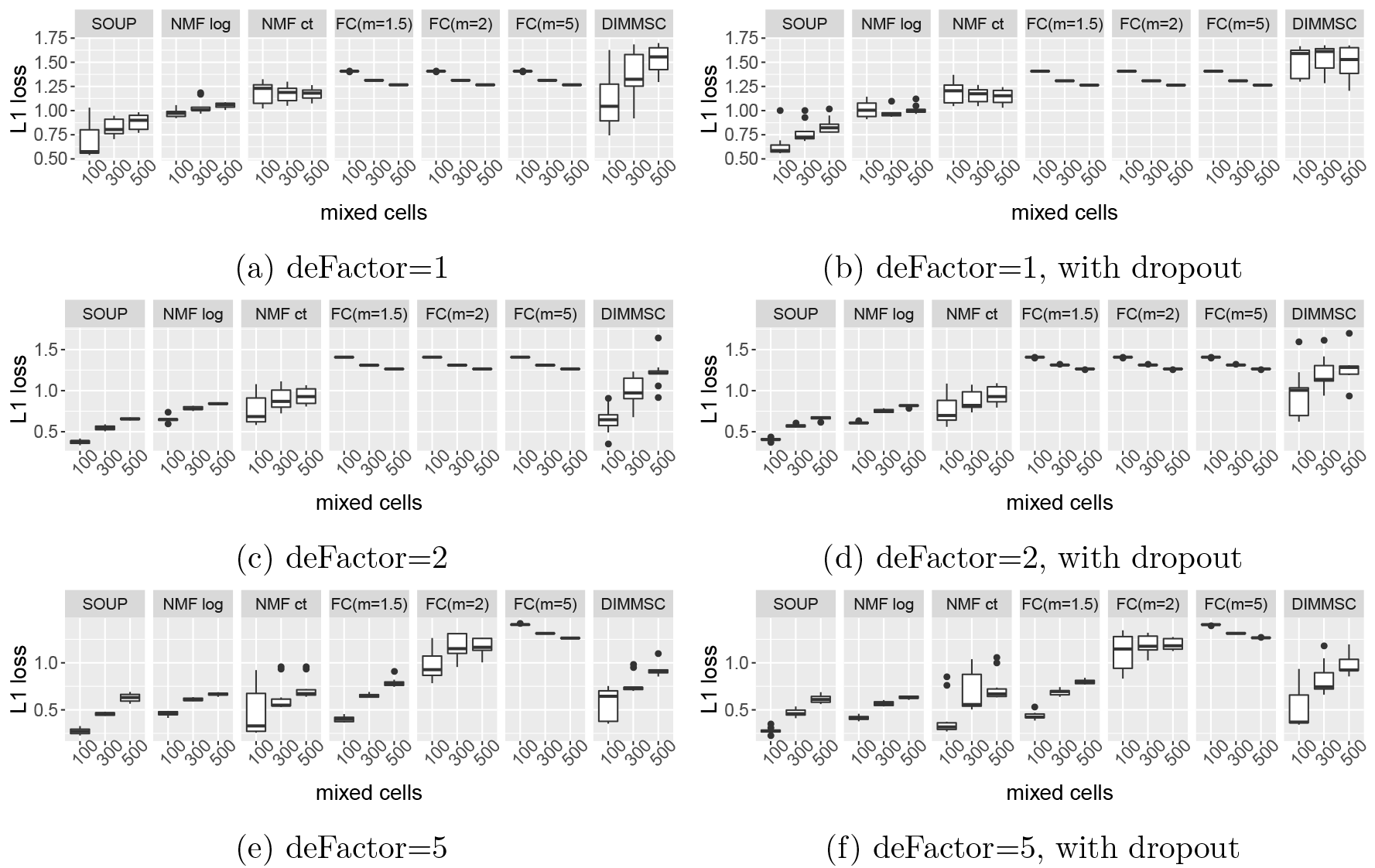
Boxplots of average *L*_1_ losses of SOUP estimated memberships in simulations with 10 repetitions, using three differential expression factors `deFactor` ∈ {1, 2, 5}, with and without dropouts. 300 pure cells are simulated from 4 clusters with probability (0.2, 0.2, 0.2, 0.4), and {100, 300, 500} mixed cells simulated along the developmental trajectory of type1 → type2 → {type3 or type 4}.

**Figure S3:**
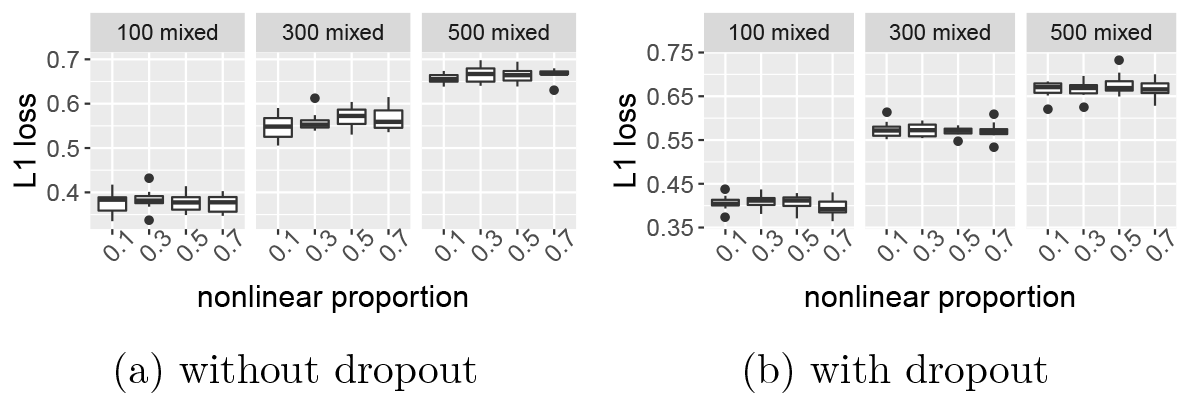
Boxplots of average *L*_1_ losses of SOUP estimated memberships in simulations with 10 repetitions. 300 pure cells are simulated from 4 clusters, and {100, 300, 500} mixed cells are simulated along the developmental trajectory of type1 → type2 → {type3 or type 4}, where {0.1, 0.3, 0.5, 0.7} of the genes follow nonlinear differentiation patterns between cell types.

**Figure S4:**
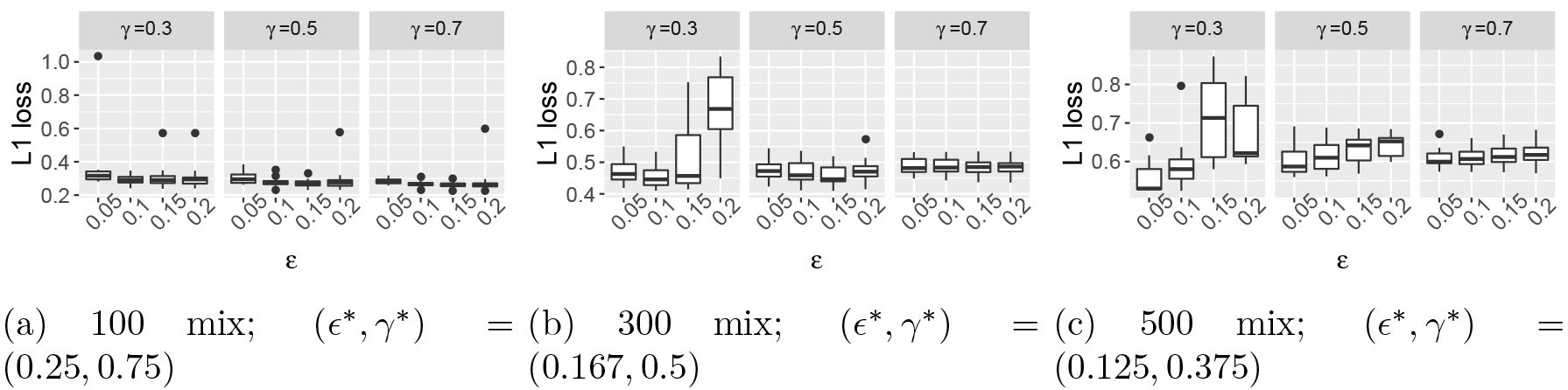
Boxplots of average *L*_1_ losses of SOUP in simulations using different tuning parameters, each repeated 10 times, where `deFactor`=5 and dropouts are included. In addition to 300 pure cells, there are {100, 300, 500} mixed cells. SOUP is applied with different tuning parameters, *ϵ* ∈ {0.05, 0.1, 0.15} and *γ* ∈ {0.3, 0.5, 0.7}. The optimal parameters (*ϵ*\*, *γ*\*) under each scenario are also listed.

### S3.5 Comparison to LOVE

One can potentially apply the LOVE algorithm (Bing et al., 2017) for *variable* clustering to *X^T^*, by treating cells as “variables”, and use the estimated allocation matrix as 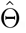. Here, we use the general routine for non-diagonal covariance matrix (corresponding to a non-diagonal *Z* matrix in our setting), and examine its performance in both simulations and real data.

We illustrate the performance of LOVE when `deFactor`=5 with 300 mixed cells. The LOVE algorithm estimates *K*, the number of clusters, as part of the procedure of finding pure cells, and requires a tuning parameter *δ* to offset the noise level in the data. In Bing et al. (2017), the authors suggest to select the optimal 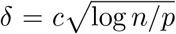 using sample splitting over a grid of *c* (note that we are treating *n* cells as variables). To gain some intuition of the effect of *δ*, we run the first step of LOVE over a grid of *c* ∈ {0.1, 0.15, …, 2.5}, and we see that the estimated number of clusters 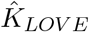 in general decreases with larger *c*, and when *c* = 2.5, LOVE underestimates the true *K* (Figure S5a). We also examine the quality of the estimated pure cells by evaluating the precision and recall rate. We repeat the simulation for 10 times, and the performance of LOVE is unstable across different runs. In addition, both the precision and recall rates of LOVE are usually lower than 0.5 (Figure S5b). On the contrary, when we vary the tuning parameter *γ* ∈ {0.1, 0.2, …, 1}, SOUP always achieves high precision in estimating the pure cells, and as soon as *γ* is larger than 0.5, the recall rate is close to 1. In addition, the variation of SOUP across different runs is also much smaller than LOVE (Figure S5c).

Next, we follow the instructions in Bing et al. (2017) to search for the optimal 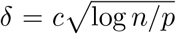 using sample splitting over *c* ∈ {0.1, 0.15, …, 2.45, 2.5}. We examine the performance with {100, 300, 500} mixed cells, with and without dropouts, each repeated 10 times. The estimated 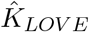 usually overestimates the true number of clusters *K* = 4 (Figure S5d). Among the three cases where 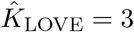, the average *L*_1_ losses of 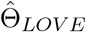 are over 40% higher than the average losses of SOUP under the same scenarios.

**Figure S5:**
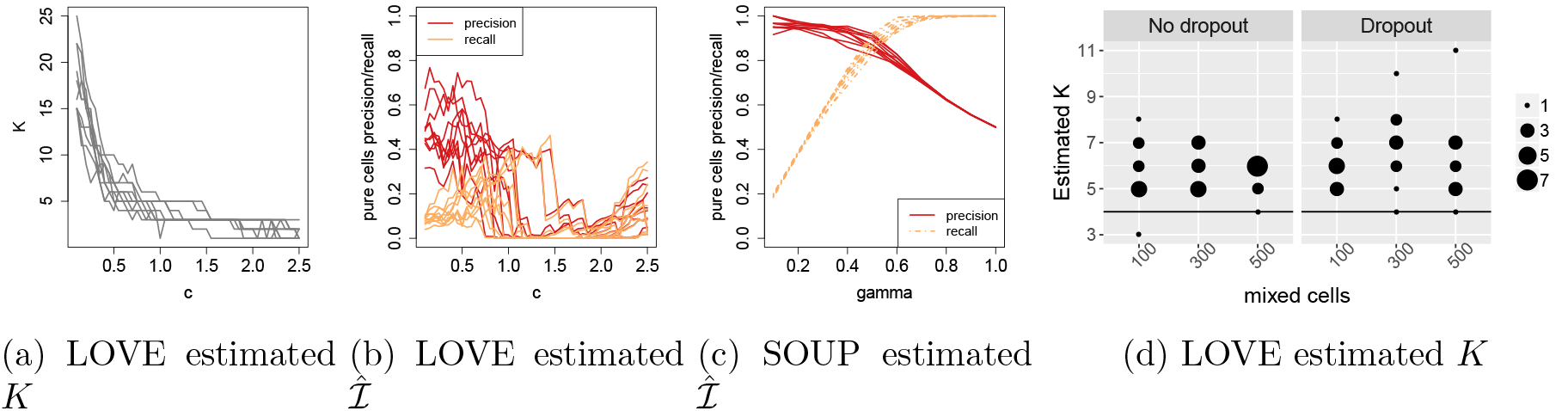
Comparison of LOVE and SOUP in 10 repetitions. We examine the scenario with and without artificial dropouts. **(a)** With 300 mixed cells, the estimated 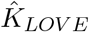 of LOVE using different tuning parameter *c*. **(b)** With 300 mixed cells, the precision and recall of LOVE estimated pure cells, using different parameter *c*. **(c)** With 300 mixed cells, the precision and recall of SOUP estimated pure cells, using different parameter *γ*. **(d)** The estimated 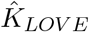 from LOVE.

Finally, we apply LOVE to the 7 public single cell datasets as in the main text. The optimal tuning parameter is selected over grid *c* ∈ {0.5, 2.1, …, 10}, where *K* can be as small as 3 and as large as over 50. Using the selected tuning parameter, the resulting 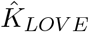 and the corresponding ARI of LOVE major types are shown in Table S7. We see that the performance of LOVE is unstable across different datasets, sometimes with substantially overestimated *K*. It achieves lower ARI when compared to SOUP, except for the Baron mouse2 dataset.

## S4 Supplementary Results

### S4.1 Discussion of fetal brain data I

The published data (Camp et al., 2015) have been normalized by Fragments Per Kilobase of transcript per Million (FPKM) and log-transformed. To apply SOUP, we first transform the data back to count scale, round to the closest integer, and then apply normalization and log transformation as usual. We start with 12,694 genes that are non-zero in at least 2 cells. After gene selection, 430 genes are retained for SOUP clustering, including 300 selected by DESCEND and 158 selected by SPCA. The original cell types are labeled according to 18 marker genes in Camp et al. (2015), of which 12 are selected by our procedure. We run SOUP with *K* = 2, 3, …, 7, and compare our hard clustering results with the published results (Figure S6a). With *K* = 5, SOUP labels are largely consistent with Camp, with two AP clusters, one BP cluster, and two neuron clusters. The sequence of different *K*’s also reveals the hierarchical structure in the data. For example, *K* = 2 separates progenitors versus neurons, and *K* = 3 gives three major groups: (i) early neural progenitors (mainly AP1), (ii) more matured progenitors and early neurons, and (iii) the matured neurons. As soon as *K* = 4, the AP2 and BP2 cells are revealed. Because BP2 cells are always clustered with early neurons, and there are always only two subtypes of neurons, we use *K* = 5 in the analysis. Notice that with *K* = 6 and *K* = 7, the effective number of major clusters is 5 and 6, respectively, meaning that there is one column of Θ that is never the largest proportion for any cell, and this situation is usually an indication of a misspecified *K*. We further visualize the soft membership of cells in the leading principal component space, and observe smooth transition from the early AP cluster to the mature neuron cluster (Figure S6b).

**Figure S6:**
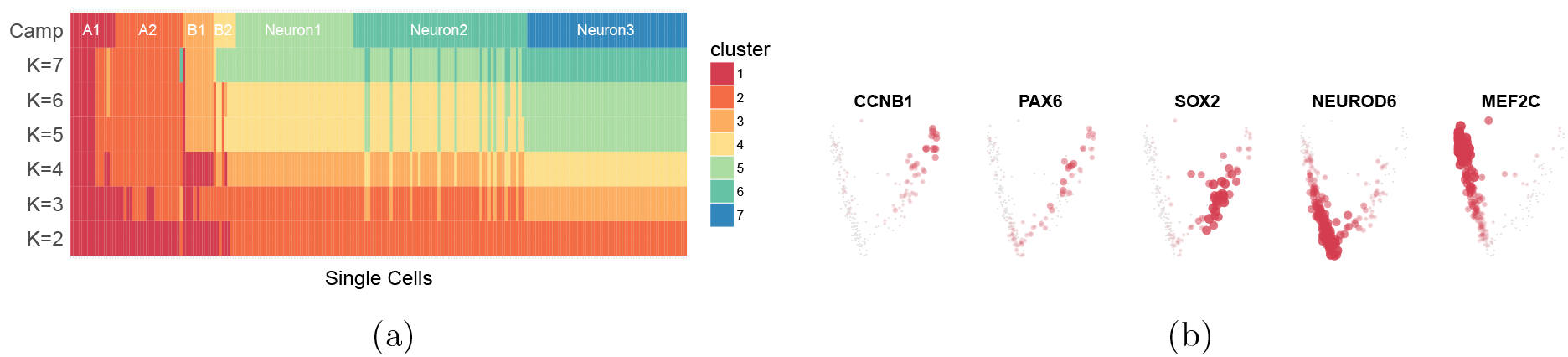
**(a)** SOUP major cluster for 220 fetal brain cells when *K* = 2, 3, …, 7, compared to Camp labels (Camp et al., 2015) on the top. Each entry is colored by the cluster assignment of each cell. **(b)** SOUP estimated soft memberships. 220 fetal brain cells are visualized in the leading 2-dimensional principal space, and each cluster is labeled by an anchor gene. In each cluster, cells with proportion ≥ 0.01 are highlighted, and the sizes and color transparency represent their estimated proportions.

We see in the main text the SOUP identified two instead of three neuron subtypes. To validate this finding, we again examine the expression levels of the 12 marker genes in the 220 fetal brain cells, where the differentiation among the three Camp neuron subtypes is ambiguous (Figure S7a).

**Figure S7:**
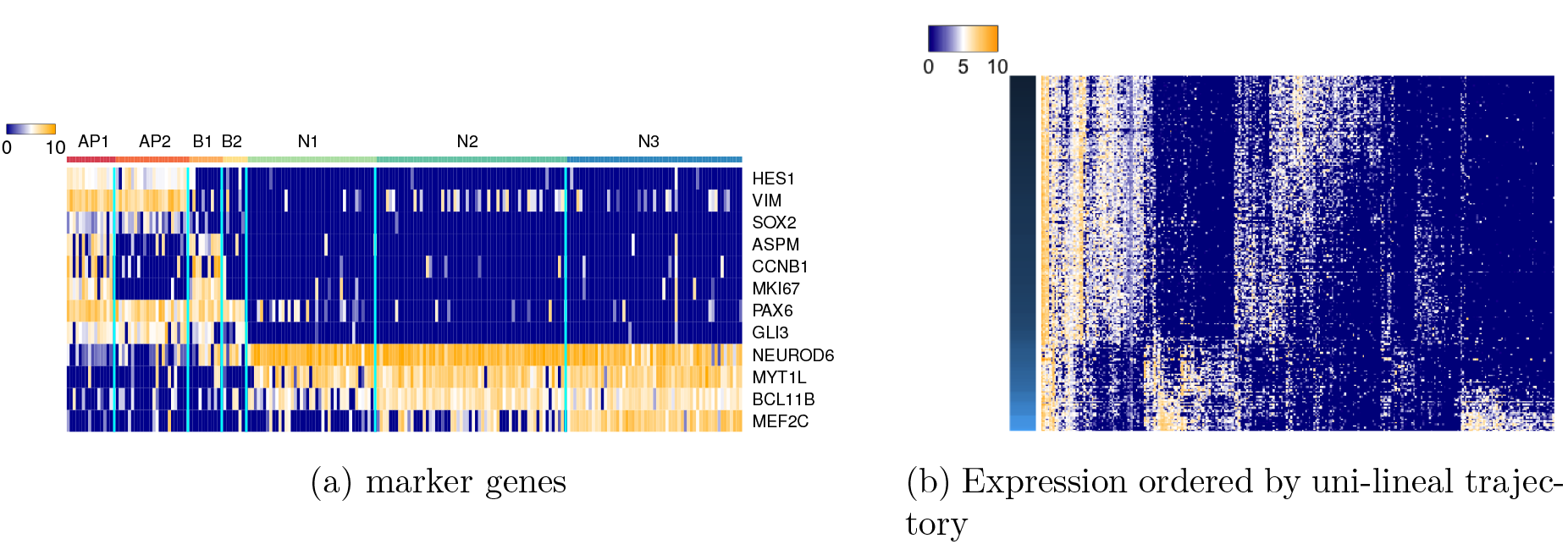
Expression levels of the 220 fetal brain cells in Camp data, visualized in log scale. **(a)** Expression levels of 12 marker genes, where cells on the columns are ordered according to Camp labels, from left to right: AP1, AP2, BP1, BP2, N1, N2, and N3. **(b)** Expression levels of the 315 PC genes identified in Camp et al. (2015), where 220 fetal brain cells on the rows are ordered according to single lineage SOUP developmental trajectory, indicated by the color bar on the left.

Next, we examine the SOUP estimated developmental trajectory. We evaluate the change of expression profiles of the 315 PC genes identified in Camp et al. (2015), along the SOUP trajectory (Figure S7b). We observe a smooth transition along different periods that is consistent with the developmental order, suggesting the SOUP estimation is sensible.

### S4.2 Discussion of fetal brain data II

We applied SOUP to the entire sample of 2,309 cells with an aim to identify the IN and MIC cells and remove them so that we can perform trajectory analysis on the cells that develop in the PFC. We rely on the Zhong labels to identify the type of cells in a cluster, but we did not want to rely on these labels exclusively, so we attempted to identify clusters largely populated by IN or MIC cells and remove them prior to our trajectory analysis. In total we run SOUP 3 times, the first two steps are required to identify and remove the IN and MIC clusters.

Step 1. The number of genes selected using Descend score *>* 8 was 627, the number of genes chosen using sparse PCA was 198 for a total of 744 distinct genes. The maximum cross validation score suggests 11 cell types; however, the best classification with at least 2 pure cells per cluster has 6 distinct cell types. Based on the results we remove cluster 3 (MIC) and cluster 4 (IN). Cluster 6 is ambiguous so we proceed with these cells included (Table S1).

**Table S1:** Contingency table of Zhong labels and major SOUP labels of all cells over the 6 clusters.
Step 2. With 1806 cells remaining, we select 616 unique genes. Cross-validation and the requirement of at least 2 pure cells per cell type led to fitting 8 distinct cell types. Based on the results we removed cluster 3 which has a large number of IN cells (Table S2).

| Zhong<br>Labels | SOUP Clusters |  |  |  |  |  | Total |
| --- | --- | --- | --- | --- | --- | --- | --- |
|  | 1 | 2 | 3 | 4 | 5 | 6 | Total |
| EN | 932 | 17 | 0 | 8 | 2 | 98 | 1057 |
| NPC | 80 | 185 | 0 | 0 | 25 | 0 | 290 |
| OPC | 7 | 24 | 0 | 1 | 73 | 12 | 117 |
| AST | 0 | 0 | 0 | 0 | 64 | 12 | 76 |
| MIC | 0 | 2 | 64 | 0 | 0 | 2 | 68 |
| IN | 40 | 1 | 0 | 430 | 4 | 226 | 701 |
| Total | 1095 | 229 | 64 | 439 | 168 | 350 | 2309 |

**Table S2:**
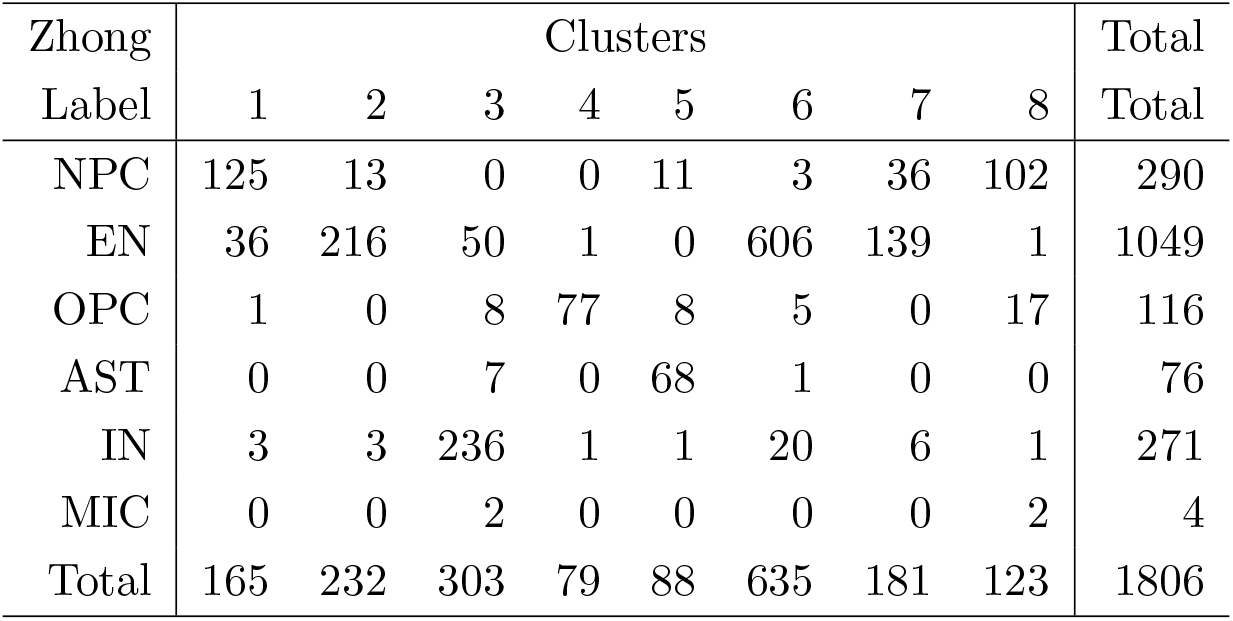
Contingency table of Zhong labels and major SOUP labels of all cells over the 8 clusters after first stage of MIC and IN clusters.
Step 3. A total of 1503 cells remain. We used 527 genes for clustering in the final step. Cross-validation and the requirement of having at least two pure cells in each cell type point towards 7 distinct cell types. The resulting clusters have very few cells that were classified as MIC or IN by Zhong and these cells are distributed across many clusters, so we proceed with our analysis assuming these cells may have been mislabeled by Zhong. The 7 clusters are labeled as OPC, AST, NPC, NPC, NPC/EN, EN and EN cells, respectively, based on the Zhong labels (Table S3).

**Table S3:**
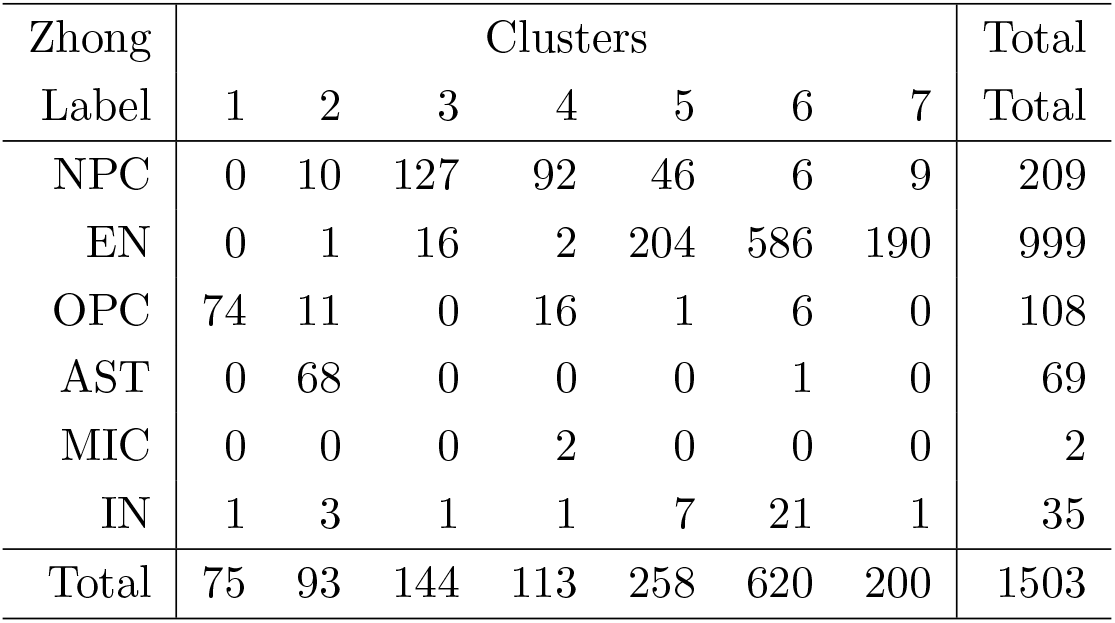
Contingency table of Zhong labels and major SOUP labels of all cells over the 7 clusters after final removal of MIC and IN clusters. Distribution of cell types over the 7 clusters assigned by hard SOUP.

| Zhong | Clusters |  |  |  |  |  |  | Total |
| --- | --- | --- | --- | --- | --- | --- | --- | --- |
| Label | 1 | 2 | 3 | 4 | 5 | 6 | 7 | Total |
| NPC | 0 | 10 | 127 | 92 | 46 | 6 | 9 | 209 |
| EN | 0 | 1 | 16 | 2 | 204 | 586 | 190 | 999 |
| OPC | 74 | 11 | 0 | 16 | 1 | 6 | 0 | 108 |
| AST | 0 | 68 | 0 | 0 | 0 | 1 | 0 | 69 |
| MIC | 0 | 0 | 0 | 2 | 0 | 0 | 0 | 2 |
| IN | 1 | 3 | 1 | 1 | 7 | 21 | 1 | 35 |
| Total | 75 | 93 | 144 | 113 | 258 | 620 | 200 | 1503 |

**Table S4:** Statistics associated with final clustering of cells, post removal of IN and MIC clusters (Table S3). Average gestational age of each of cell type cluster and average maximum theta for the primary type.

| Cluster | Type | Age | maxP |
| --- | --- | --- | --- |
| 1 | OPC | 23.1 | .77 |
| 2 | AST | 24.1 | .75 |
| 3 | NPC | 11.7 | .51 |
| 4 | NPC | 16.0 | .73 |
| 5 | EN/NPC | 15.8 | .52 |
| 6 | EN | 18.8 | .60 |
| 7 | EN | 10.3 | .71 |

The majority membership probabilities suggest that many cells are in the transitional phase (Figure S8). Trajectory analysis will facilitate our investigation of these transitional cells.

Cluster 4 was chosen to represent the earliest cell type in the lineage analysis because it contains the fewest cells classified by Zhong as EN. Based on 3 dimensions in Slingshot (nPC = 3), starting with cluster 4, we identified two lineages (Figure 7). One neuronal lineage, L1: 4 → 3 → 7 → 6 → 5 moves from NPC cells to various stages of EN cells; another glia cell lineage, L2: 4 → 3 → 1 → 2 moves from NPCs to AST cells.

Next, we examined how sensitive SOUP is to the number of genes used. We chose genes using the DESCEND algorithm with a less stringent cutoff so that the total number of genes selected is 1917 rather than 527 for the analysis of the 1503 cells selected above. Using more genes we chose *K* = 6 clusters (1’,…,6’) and the clusters are surprisingly similar to the 7 discovered with fewer genes (1,…,7), with the exceprtion of cluster 6’ (Table S5). Using more genes we fail to differentiate clusters 6 and 7 that contain EN cells. Moreover, half of cluster 5 cells migrate to cluster 6’. Clusters 5, 6 and 7 are the most similar of all the clusters and 15-20% of the elements clustered to these cell types have a substantial probability of being in transition between these clusters (based on the estimated parameters). We conclude that SOUP is not highly sensitive to the number of informative genes chosen. Nevertheless, it appears that a stricter list of genes can separate clusters slightly better.

**Figure S8:**
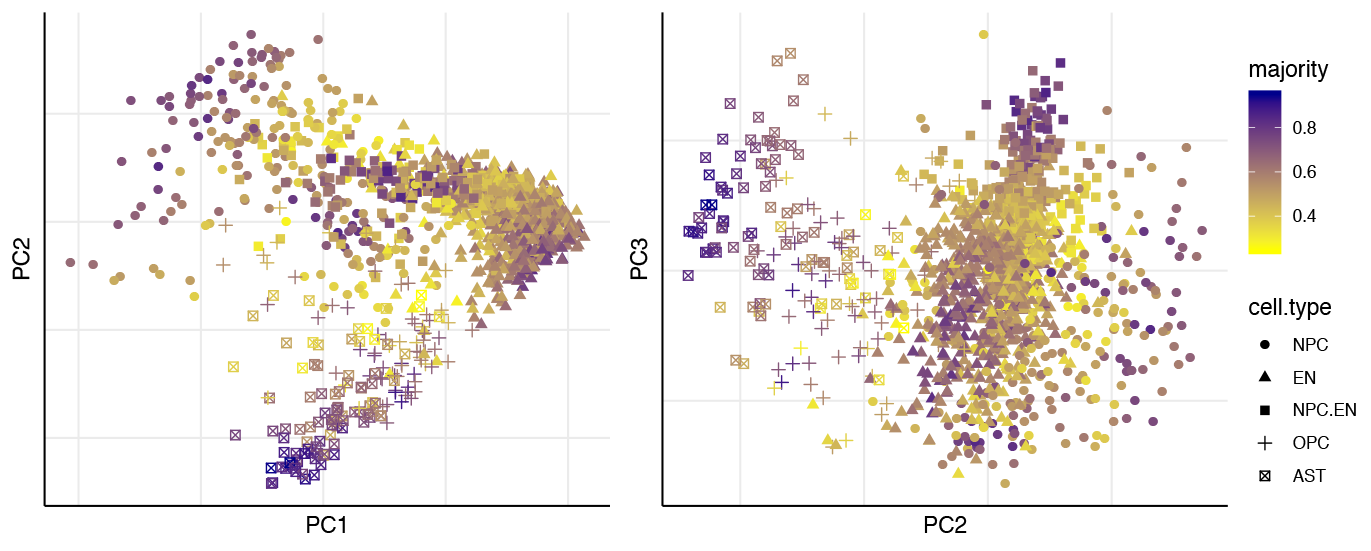
SOUP estimated majority membership probabilities of all cells in Zhong dataset, excluding IN and MIC.

**Table S5:** Contingency table of SOUP labels calculated using 527 genes versus 1917 genes.

| 527<br>Genes | 1917 Gene Clusters |  |  |  |  |  | Total |
| --- | --- | --- | --- | --- | --- | --- | --- |
|  | 1' | 2' | 3' | 4' | 5' | 6' | Total |
| 1 | 62 | 2 | 0 | 0 | 3 | 1 | 67 |
| 2 | 0 | 88 | 0 | 2 | 0 | 0 | 90 |
| 3 | 0 | 5 | 4 | 103 | 13 | 15 | 140 |
| 4 | 1 | 0 | 4 | 3 | 102 | 1 | 111 |
| 5 | 0 | 3 | 108 | 1 | 5 | 114 | 231 |
| 6 | 0 | 3 | 0 | 3 | 0 | 553 | 559 |
| 7 | 0 | 0 | 5 | 1 | 1 | 191 | 198 |
| Total | 63 | 100 | 121 | 113 | 124 | 875 | 1396 |

A key advantage of SOUP is its soft membership. For example, we can use the probability of membership SOUP yields to explore the molecular features inherent in the development of cell types. Here we explore the development of neurons in clusters 5 and 6, the last two clusters in the neuronal lineage in Figure 7. We select these two clusters for their features. Cluster 6 falls tightly along the neuronal trajectory and is composed, predominantly, of cells labeled neurons (EN) by Zhong. Alternatively, cluster 5, which falls at the end of the trajectory, shows greater spread, curls back toward NPC cells, and 17.8% of its cells are labeled NPC by Zhong. To elicit features of neuronal development, we contrast cells in a cluster that either fall near the trajectory, but far from the soft cluster center (Group 1), or cells in the cluster that fall far from the trajectory and the soft cluster center (Group 2). Typically, membership to the cluster is less certain for cells away from the cluster center and we enforce this feature by only analyzing cells with membership probability < 0.5. Label Group 1’s less mature cells – determined by the trajectory– as set **A** and more mature cells as set **B**; for Group 2, less mature cells fall in set **C** and more mature cells fall in set **D** (Figure 7a). The idea is that the contrast of gene expression patterns of **A** versus **B** informs about which proteins are changing as neurons of this trajectory develop, while the contrast of **C** versus **D** and **B** versus **D** could yield information about development of alternative types of neurons. This latter analysis is motivated by the fact that there is a wide diversity of types of excitatory neurons in the human brain.

For cluster 6, 10, 43, 26 and 10 cells fall into the **A** through **D** sets, respectively (Figure S9a). When the mean expression of 527 genes in cells from **A** versus **B** is contrasted (Table S8), 135 genes show significant changes in expression (p ¡ 0.05, uncorrected, t-test): 38.5% of the genes marking ENs (5/13), according to Zhong, and 14.3% of their NPC markers (24/168). Mean expression for all five neuronal markers increases from **A** to **B**, consistent with maturing neurons, and the enrichment for neuronal markers among differentially expressed genes is 3.7 (*p* = 0.038). Tellingly, one of the NPC markers is *MYT1L*, which, while first expressed in NPCs, is also expressed in neurons, where it locks in and is essential for the maintenance of neuronal state Mall et al. (2017). *MYT1L* also increases in expression from **A** to **B** cells. Many of the 24 differential NPC markers do too (Table S8), although some of these genes are also critical for neurons (e.g., *NRXN1*, encoding a synaptic protein). The contrast of mean gene expression for **C** versus **D** cells reveals remarkably similar patterns to those seen in **A** versus **B**, again consistent with developing neurons (Table S8, Figure S9b). Given that **B** and **D** show the hallmarks of neurons, we next evaluated these cell clusters in detail for all 13 neuronal markers identified by Zhong and including *MYT1L*. Of these, one show significant differential mean expression for **B** versus **D**, *IGFBPL1*, p = 0.0023. Notably, most of the 13 neuronal markers are elevated in **B** versus **A** (13/13) and **D** versus **C** (9/13) (Table S8). Together these data reveal several features of the cells in cluster 6: the cells show expression patterns consistent with immature neurons early in the trajectory of this lineage and maturing neurons later in the lineage; the level of expression of *MYT1L* is consistent with commitment to neuronal fate in maturing neurons; and the differences in gene expression between cells in cluster **B** versus **D** is suggestive the cells could differentiate into different neuronal cell types.

For cluster 5, 10, 8, 7 and 15 cells fall into the **A** through **D** sets, respectively (Figure S9a). When the mean expression of genes in cells from **A** versus **B** are contrasted (Table S8), 164 genes show significant changes in expression: 15.4% of the genes marking ENs (2/13) and 44.6% NPC markers (75/168). Curiously, the expression of the differential neuronal markers diminishes slightly from **B** to **A** (2/2), whereas most of the NPC markers increase (70/74), and the enrichment of markers favors NPCs, not neurons (Odds ratio = 0.23, *p* = 0.045). Again, the patterns seen for **A** versus **B** are similar to those for **C** versus **D** (Table S8). Because the average expression of NPC markers increases in cells found later in the trajectory, it is reasonable to ask if **B** and **D** cells of cluster 5 are truly immature neurons? Assessing neuronal markers and *MYT1L*, it is clear that the sets of cells (**A**-**D**) exhibit the hallmarks of maturing neurons (Figure S9c), which can also be seen by comparing the patterns of gene expression for cluster 6 versus cluster 5 (Figure S9a vs. b). Moreover, when we sub-cluster the cells in cluster 5 separately, we obtain four clusters, two of which we will examine more closely. One corresponds almost perfectly to Zhong’s NPC cells and the other to a large group to Zhong’s neurons, and these groups of cells are indistinguishable for mean expression of neuronal markers and *MYT1L* (data not shown). At least two possibilities spring from these observations: either the developmental patterning of gene expression in the neurons of cluster 5, which are likely deep-layer projection neurons according to their GW age, are not as well understood as the literature suggests; or there is some contamination of cluster 5 neurons with NPC cells, for instance, some pairs of different cell types sequenced together. The former is hard to evaluate. If the latter were true, we would expect a greater diversity of genes to be expressed in cluster 5 cells. This pattern is observed in the data (Figure S9d) and indeed cluster 5 cells express significantly more genes than any other cluster: the most similar cluster is 4, which are NPC cells (two-sided *p* = .043); for neurons, cluster 7 is most similar, but it is highly significantly different (*p* < 10^−16^). Thus, the cells from cluster 5 show strong and consistent expression of neuronal markers regardless of location in the trajectory and cluster space (Figure S9a,c), suggesting that many or most of these cells are maturing neurons; however, there is also some contradictory evidence because mean expression of a substantial fraction of NPC markers increases in mean expression from early to late in the developmental trajectory.

**Figure S9:**
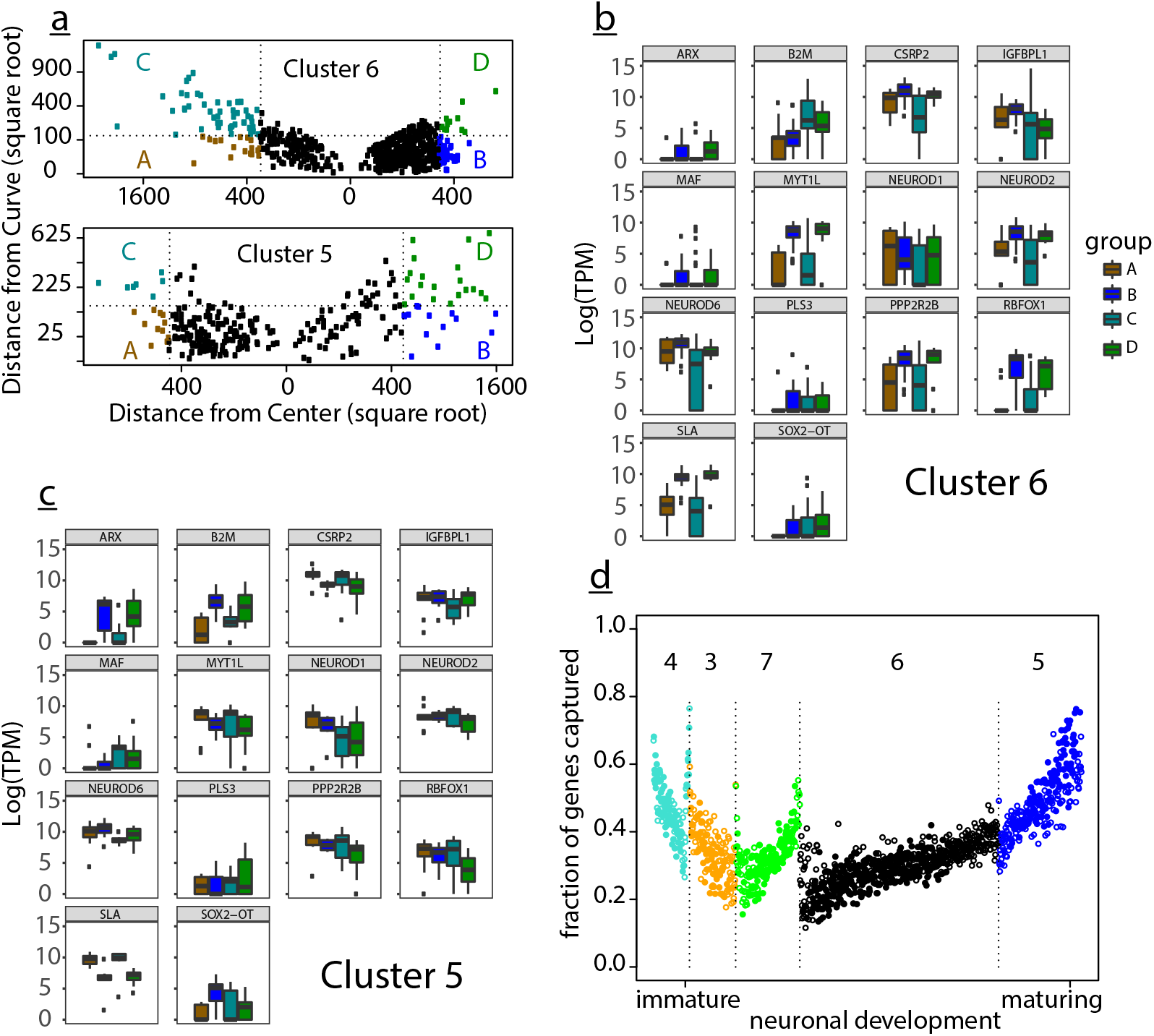
Selection and gene expression patterns in neuronal lineage cells from cells in clusters 5 and 6 (Figure 7, main). **(a)** Identifying four sets of cells: A & B, cells far from the cluster center but near the neuronal developmental trajectory (curve), where A is earlier in the projected lineage and B is later; C & D, cells far from the cluster center and far from the trajectory, where C is earlier in the projected lineage and D is later. Far is defined on the distribution of distances in three dimensions in the principal component space, and specifically ≥ 20% from the center of the cluster and membership *θ* < 0.5; near is defined likewise, as < 20% from the curve. **(b)** For sets A to D of cluster 6, expression of the 13 genes cited as neuronal markers by Zhong et al. (2018) and one cited as NPC, *MYT1L*, which is critical for determination and maintenance of neuronal state. Expression in log of transcripts per million (TPM). **(c)** For sets A to D of cluster 5, expression of neuronal markers and *MYT1L*. Expression is scaled in log of transcripts per million (TPM). **(d)** Number of genes with expression greater than zero for each cell in the neuronal lineage, by cluster.

## S5 Supporting Tables

**Table S6:** Major characteristics of 7 public datasets, including human brain cells from Darmanis et al. (2015), and pancreatic cells from four human donors and two mouse strains from Baron et al. (2016). For SOUP, we list the number of selected genes by SPCA and DESCEND, the ARI using two different *γ* ∈ {0.5, 0.8}, as well as the total runtime in seconds, including gene selection and clustering, where DESCEND is paralleled with 5 cores, and the computation is conducted on a linux computer equipped with AMD Opteron(tm) Processor 6320 @ 2.8 GHz.

|  | Dataset characteristics |  |  | SOUP |  |  |  |
| --- | --- | --- | --- | --- | --- | --- | --- |
| | K | # cells | # genes | # sel. genes | runtime | ARI ($\gamma = 0.5$) | ARI ($\gamma = 0.8$) |
| Darmanis | 8 | 420 | 22,085 | 1,106 | 40 | 0.8890 | 0.9071 |
| Baron human1 | 14 | 1,937 | 20,125 | 1,632 | 227 | 0.7513 | 0.7519 |
| Baron human2 | 14 | 1,724 | 20,125 | 1,571 | 188 | 0.7946 | 0.7502 |
| Baron human3 | 14 | 3,605 | 20,125 | 1,923 | 848 | 0.4071 | 0.5030 |
| Baron human4 | 14 | 1,303 | 20,125 | 1,516 | 112 | 0.7756 | 0.5092 |
| Baron mouse1 | 13 | 822 | 14,878 | 895 | 50 | 0.5782 | 0.4541 |
| Baron mouse2 | 13 | 1,064 | 14,878 | 1,324 | 78 | 0.6280 | 0.4093 |

**Table S7:** Selected *K* and corresponding SOUP ARI on 7 public datasets (Baron et al., 2016; Darmanis et al., 2015). We select the optimal *K* among {2, …, 20}, using (i) optimal Calinski-Harabasz (CH) index or Silhouette score, computed from K-means hard-clustering results of the pure cells, where pure cells are identified by SOUP with *γ* = 0.8; and (ii) minimal sample splitting (SS) prediction error of SOUP, averaged over 10 repetitions. We further compare the ARI of SOUP hard assignments using different selected *K*’s. Finally, we present the estimated *K* from LOVE, where the tuning parameter is selected by cross validation over grid *c* ∈ {0.5, 0.6, …, 9.9, 10}, and the number of cross validation is set to the default choice of 50. All algorithms are applied to the selected genes by DESCEND and SPCA.

| Dataset | Selected $K$ | | | | SOUP ARI with selected $K$ | | | LOVE | |
| --- | --- | --- | --- | --- | --- | --- | --- | --- | --- |
| | Reference | CH | Silhouette | SS | CH | Silhouette | SS | $\hat{K}$ | ARI |
| Darmanis | 8 | 3 | 2 | 6 | 0.5073 | 0.1844 | 0.8957 | 27 | 0.4606 |
| Baron human1 | 14 | 2 | 4 | 13 | 0.3433 | 0.3908 | 0.7512 | 18 | 0.6796 |
| Baron human2 | 14 | 2 | 3 | 10 | 0.3329 | 0.3652 | 0.8986 | 13 | 0.6322 |
| Baron human3 | 14 | 2 | 2 | 11 | 0.3944 | 0.3944 | 0.6034 | 51 | 0.2068 |
| Baron human4 | 14 | 2 | 2 | 5 | 0.3439 | 0.3439 | 0.7509 | 11 | 0.6846 |
| Baron mouse1 | 13 | 2 | 2 | 7 | 0.5411 | 0.5411 | 0.8533 | 14 | 0.5158 |
| Baron mouse2 | 13 | 2 | 2 | 7 | 0.2823 | 0.2823 | 0.6392 | 16 | 0.8602 |

Table S8: Attached as supplementary material.

